# Neurons entrain and shape an intrinsic arteriole rhythm that distributes blood across the cortical mantle

**DOI:** 10.64898/2026.11.05.764641

**Authors:** Jacob Duckworth, Thomas Broggini, Karishma Chhabria, Philip Neill, María Sancho, Marie Sellier-Prono, Satish Iyengar, Mark T. Nelson, Massimo Vergassola, David Kleinfeld

## Abstract

The cerebral arteriole network distributes blood across cortex through branches whose diameters oscillate near a frequency f_vaso_ ∼ 0.1 Hz. We asked whether the spatiotemporal pattern of these vaso-oscillations reflects neuronal entrainment of an intrinsic vascular oscillation. Three findings support this: First, suppressing endothelial hyperpolarization left resting-state oscillations intact but weakened the vascular response to neural activity, showing that vasomotion originates within the arteriole network rather than following neuronal drive. Second, arterioles amplify sensory drive selectively rather than faithfully: neural activity near 0.1 Hz is resonantly enhanced to form a strong peak in the vascular response. Third, neuronal frequency sets spatial reach, and areas compete: when two neighboring cortical areas are driven simultaneously, the one nearer f_vaso_ captures more vascular territory. Neurons therefore entrain an intrinsic vascular rhythm and stimulation frequency governs how vasomotion distributes blood, shapes functional brain images, and may influence interstitial fluid transport.

## INTRODUCTION

The cerebral vasculature is a sensorimotor organ. It is both the thoroughfare for the flow of blood yet adjusts the flow to accommodate regional metabolic demands via regulation of pial and parenchymal arterioles that span the cortical mantle^1^. This process, denoted functional hyperemia, occurs over periods from one to tens of seconds and is sufficiently stereotyped such that imaging methods including functional magnetic resonance imaging (fMRI) and functional near infrared spectroscopy (fNIRS) can be used to noninvasively infer neural activity from spectroscopic changes in blood flow and oxygenation^2,3^. As a description of the relation of flow to neural activity, linear response theory was used extensively in early studies^4,5^. Yet subsequent work showed it to be less accurate when applied across brain regions and variable stimuli conditions^6–9^. Pertinently, arteriole smooth muscle cells (SMCs) exhibit spontaneous activity that drives cycles of dilation and constriction at ∼ 0.1 Hz in a process termed vasomotion^10–12^. Although hemodynamic metrics used to study the brain’s functional networks are dominated by activity near 0.1 Hz^13–15^, the influence of intrinsic vasomotion on the spatiotemporal hemodynamic representation of neural activity remains unexplored.

Vasomotion has been investigated extensively in human and animal vascular beds^10,11^. However identifying the driving force underlying vaso-oscillations is historically difficult, as systemic influences can modulate blood flow near the 0.1 Hz frequency band of spontaneous SMC activity^16,17^. Vasomotion occurs in isolated arterioles, suggesting a local drive, albeit often requiring initiation with norepinephrine or K^+^ channel blockers, or the use of arterioles from hypertensive animals^11,18–20^. *In vivo* studies further support local drive, as oscillation frequencies vary spatially within individual tissues^10,21^, and vasomotion persists in the absence of systemic ∼ 0.1 Hz blood pressure oscillations, i.e., Mayer waves^22,23^. In the brain, vasomotion entrains to the envelope of gamma-band neural activity^24^, and although questions remain on the independence of resting-state cerebral vasomotion from neural drive, preliminary experiments indicate that low frequency hemodynamic variability remains after broad neural blockade^25^. Based on existing evidence that vasomotor activity can phase lock or unlock to systemic and neural influences, we hypothesized that the pial arterioles behave as oscillators whose amplitude and phase are determined by the spectral content of inputs relative to the intrinsic vasomotor frequency.

Cerebral arterioles receive diverse input, including from cortical neurons that initiate functional hyperemia on the seconds time scale^1^. These functional responses can be regulated by neuromodulatory activity^26^, and appear robust to the loss of any single neurovascular messenger^27^. A common pathway is the integration and relay of parenchymal vascular hyperpolarization to the pial arterioles; in fact, individual arteriole dilations correlate more strongly with downstream arbor neuronal activity than proximal activity^28^. Central to this process is the potassium inward rectifier channel K_IR_2.1 (Kcnj2) which is expressed in capillary and arteriole endothelial cells^29,30^. Potassium efflux during neural activity activates these channels, causing membrane hyperpolarization that activates neighboring K_IR_2.1 channels^31^ and, in parallel with slower mechanisms^32,33^, mediates rapid neurovascular signaling (∼ 2 mm/s)^34,35^.

Here we assess the dynamics of the pial arteriole network in awake mice at the resolution of individual arterioles across the cortical mantle. We ask: (1) What is the spectral character of pial arterioles that oscillate at rest and how does it vary between cortical regions? Notably, arterioles with narrow spectra, i.e., nearly sinusoidal variations in diameter, would be unreceptive to broadband neural input and prone to potentially deleterious constrictions^36^. (2) Is the rhythmicity of resting-state vaso-oscillations an intrinsic property of arterioles *in vivo*, or is it dependent on neurovascular coupling? Notably, in the absence of pharmacological manipulations, arteriole preparations outside of the normal brain milieu rarely oscillate^18^. (3) When do nonlinearities in the arteriole response lead to qualitative changes in arteriole vasodynamics? The time series from MR images are commonly modeled by a convolution of neural input with a hemodynamic response function (HRF). This assumes linearity and time-invariance^4,37,38^, as occur in anesthetized and thus weakly responding animals^39^. Yet nonlinear effects are well documented in the BOLD response^7,9^. (4) Do pial arterioles undergo significant resonance enhancement when driven near their vasomotor frequency? Resonance is a generic property of all driven oscillatory systems and is thus an expected albeit yet undescribed aspect of brain arteriole dynamics. Resonance should impact the signal strength of MR imaging. (5) Does resonance impact the spatial extent as well as the amplitude of vaso-oscillations? Notably, does spectral content as opposed to physical proximity dominate the patterns of vasomotor activity when neighboring cortical regions are excited? (6) Can subcortical neural activity associated with homeostasis entrain pial arterioles when driven similarly to sensory systems? We focus on the rostral ventrolateral medulla (RVLM), a hypoxia-sensitive vasoactive nucleus^40^, that elicits cerebral blood flow increases upon activation^41^.

## RESULTS

Measurements were performed using either widefield one-photon imaging or two-photon laser-scanning imaging in awake, head-fixed mice. The field of view spanned both hemispheres of the cortical mantle (**Figure 1A,B**). We used a thinned skull preparation to gain optical access while minimizing mechanical deformation of the brain and potential physiological changes caused by inflammation^42,43^. Fluorescence was collected from the genetically encoded calcium indicators GCaMP8.1 expressed in SMCs^44^ and jRGECO1a expressed in excitatory neurons via the Thy1 promoter^45^. Previous two-photon imaging experiments showed that pial GCaMP8.1 fluorescence is highly correlated with arteriole diameter, enabling the use of this signal as a distributed measure of diameter in the widefield system^46^. We make spectrally separate measurements of arteriole and excitatory neuronal activity by alternately exciting GCaMP8.1 and jRGECO1a at a sampling rate of 10 Hz (**Figure 1C**).

**Figure 1:**
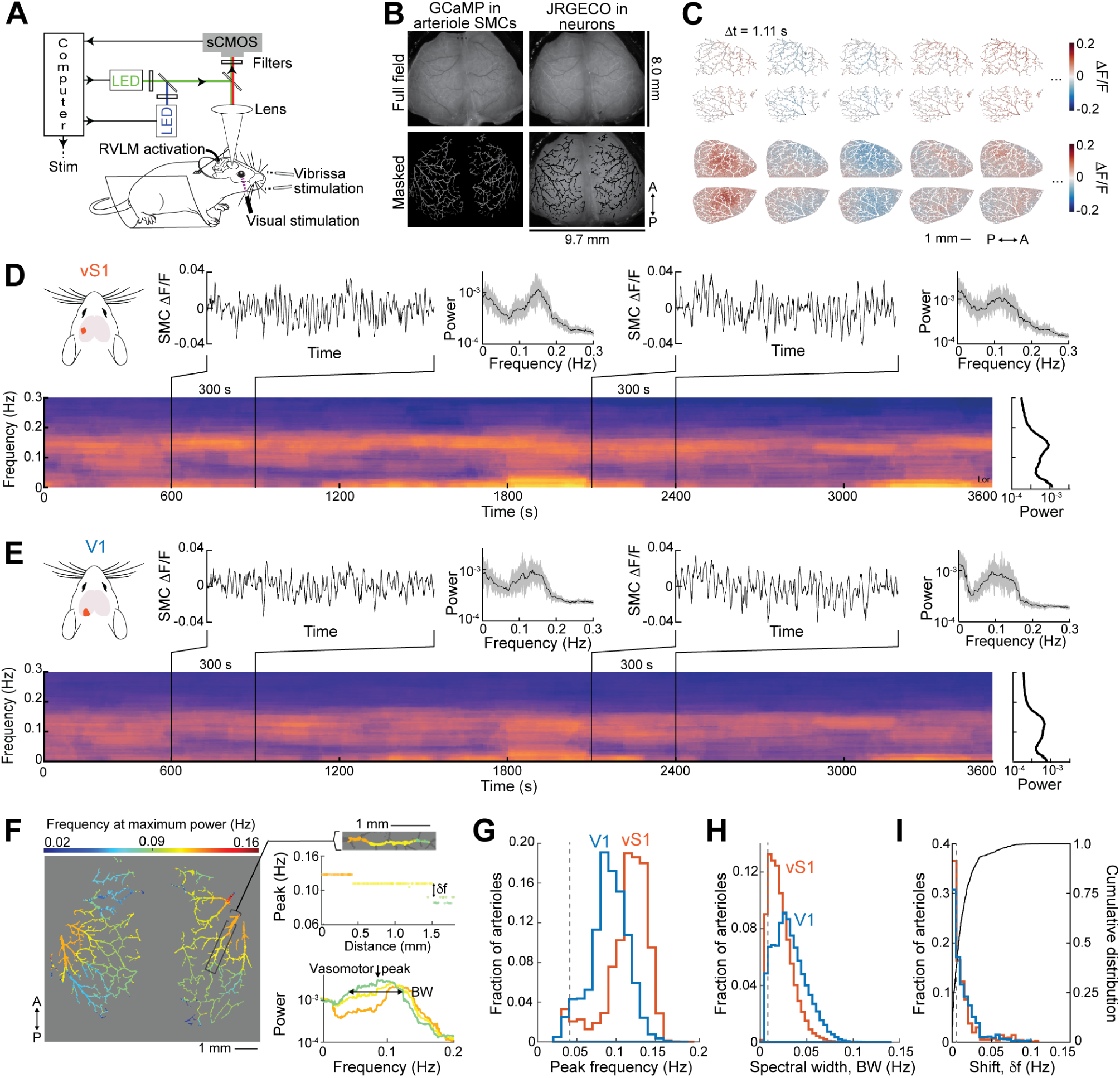
Spatiotemporal characteristics of resting-state pial arteriole activity. **(A)** Schematic of the dual-channel wide-field imaging setup. A Prime95B sCMOS camera with interleaved LED illumination and bandpass filters capturing GCaMP fluorescence in arteriole smooth muscle cells (SMCs) and jRGECO fluorescence in neurons simultaneously through a transcranial window, with concurrent stimulation options (RVLM activation, vibrissa stimulation, and visual stimulation). **(B)** Representative full-field (top) and vessel-masked (bottom) fluorescence images of GCaMP in arteriole SMCs (left) and jRGECO in neurons (right), showing the cortical vascular map across a ∼9.7 × 8.0 mm field of view; P, posterior; A, anterior. **(C)** Spatiotemporal timeseries of fractional fluorescence changes (ΔF/F) across the dorsal cortex at sequential timepoints (Δt = 1.11 s), displaying spontaneous vasomotor (top, GCaMP) and neural (bottom, jRGECO) activity. Scale bar, 1 mm; orientation: P, posterior; A, anterior. **(D)** Vasomotor dynamics recorded from summed arterioles in primary somatosensory (vS1) cortex. Example SMC ΔF/F traces (300 s epochs at 600 s and 2100 s) with corresponding power spectra (right insets, mean ± 95% CI) are shown alongside a time-frequency spectrogram of the full recording session (∼ 3600 s), with the mean power spectrum at far right. **(E)** Vasomotor dynamics recorded from summed arterioles in primary visual (V1) cortex, plotted as in (D). **(F)** Spatial map of peak vasomotor frequency across the cortical vasculature, color-coded from 0.02 to 0.16 Hz. Insets show peak frequency reduction as a function of distance between vS1 and V1 cortices along an example vessel segment of 2 mm (top) and the power spectrum of the individual segments with spectral width (*BW*) and vasomotor peak indicated for V1 cortex (bottom). **(G)** Peak frequency differences across experiments between arteries in the primary somatosensory cortex (vS1, red) and primary visual cortex (V1, blue), with dashed black line indicating the analysis full bandwidth. **(H)** Differences in vasomotor width across experiments between arteries in the primary somatosensory cortex (vS1, red) and primary visual cortex (V1, blue), with dashed line indicating the analysis full bandwidth. **(I)** Peak frequency discontinuity across experiments for arteries in the primary somatosensory cortex (vS1, red) and primary visual cortex (V1, blue), with black line showing the combined cumulative distribution and dashed black line indicating the analysis full bandwidth.

Sensory stimulation involved periodic puffs of air to the vibrissae and/or photic stimulation using periodic full-field illumination or drifting gratings to the eyes. In additional experiments, we used transgenic mice that expressed the red-shifted excitatory opsin ChRmine^47^ in adrenergic neurons within the brainstem region RVLM to drive subcortical activation of the arterioles across the cortical mantle. See **STAR METHODS** for details.

### Pial arterioles exhibit persistent resting-state oscillations that vary systematically across the mantle

We first characterized changes in pial arteriole diameter in the absence of external stimulation. During a set of 60-minute resting-state experiments, we observed a persistent peak in the spectral power density near 0.1 Hz, consistent with intrinsic vasomotion^10,11^ (**Figure 1D,E**). The peak in vasomotor activity varied between cortical regions, e.g., vibrissa and visual primary sensory cortices. Yet the vasomotor peak is stable across many cycles, taken as 300 to 500 s in our analysis, indicating that for these periods of time the width of the spectral peak is caused by cycle-to-cycle variability in the oscillation itself as opposed to a slow drift of an oscillator with a sharply defined frequency. The frequency with the maximum power varied systematically across the pial network, and nearby arterioles tend to group into frequency-locked regions (**Figure 1F**).

As an average across arterioles in all trials and animals, the peak vasomotion frequency is f_vaso_ = 0.11 ± 0.03 Hz (mean ± SD) in primary vibrissa sensory (vS1) cortex and f_vaso_ = 0.09 ± 0.02 Hz (mean ± SD) in primary visual (V1) cortex (**Figure 1G**; 95 experiments in 16 mice). The peak frequency observed for vS1 cortex is significantly higher than for V1 cortex (**Figure S1A**, p = 3.3×10^-17^ one-tail Wilcoxon signed rank test). More so, the vasomotor peak full width at half maximum, *BW*, is significantly narrower in vS1 than in V1 cortex (**Figures 1H** and **S1B**; median values *BW* = 0.02 and 0.03 Hz, respectively; p = 1.2×10^-13^, one-tail Wilcoxon signed rank test). Of importance in interpreting the spectral shift across the mantle, the vasomotor width (Figure 1H) is large compared to the difference in frequency (**Figure 1I**), *δf,* of 0.02 Hz in 79 % of arterioles between vS1 and V1 cortices. Thus the shift in vasomotor frequency across the cortical mantle appears to be continuous (**Figure 1F)**.

Classical resonators are assigned a quality factor^48^, defined by Q ≡ f_vaso_/BW. For data taken under resting-state conditions, we found that Q = 7.7 ± 5.6 (mean ± SD) for arterioles in vS1 cortex versus Q = 4.5 ± 4.1 (mean ± SD) for arterioles in V1 cortex (**Figure S1C**). Thus vaso-oscillators in vS1 cortex store more energy. Lastly, the power at the vasomotor peak is higher in vS1 than in V1 cortex (**Figure S1D**, p = 7.3×10^-14^ one-tail Wilcoxon signed rank test), consistent with the greater Q for this area.

### Endothelial K_IR_2.1 knockout sharpens resting state vasomotion

From a mechanistic perspective, an essential issue is if the underlying vaso-oscillations have a component intrinsic to arterioles or are solely dependent upon neurovascular coupling. The fast, ionic component of neurovascular coupling depends on the potassium inward rectifier channel K_IR_2.1^34^. Membrane repolarization is likely supported by hyperpolarization-activated cyclic nucleotide-gated (HCN) channels, which are also expressed in cerebrovascular endothelial cells^29,30^. Motivated by pioneering studies on the role of K_IR_2.1 in anesthetized mice^34^, we first investigated arteriole variability in awake mice before and after *in vivo* disconnection from neural activity, using inducible genetic knockout of endothelial cell (EC) K_IR_2.1 and administration of the HCN channel blocker ivabradine (**Figure 2A**).

**Figure 2:**
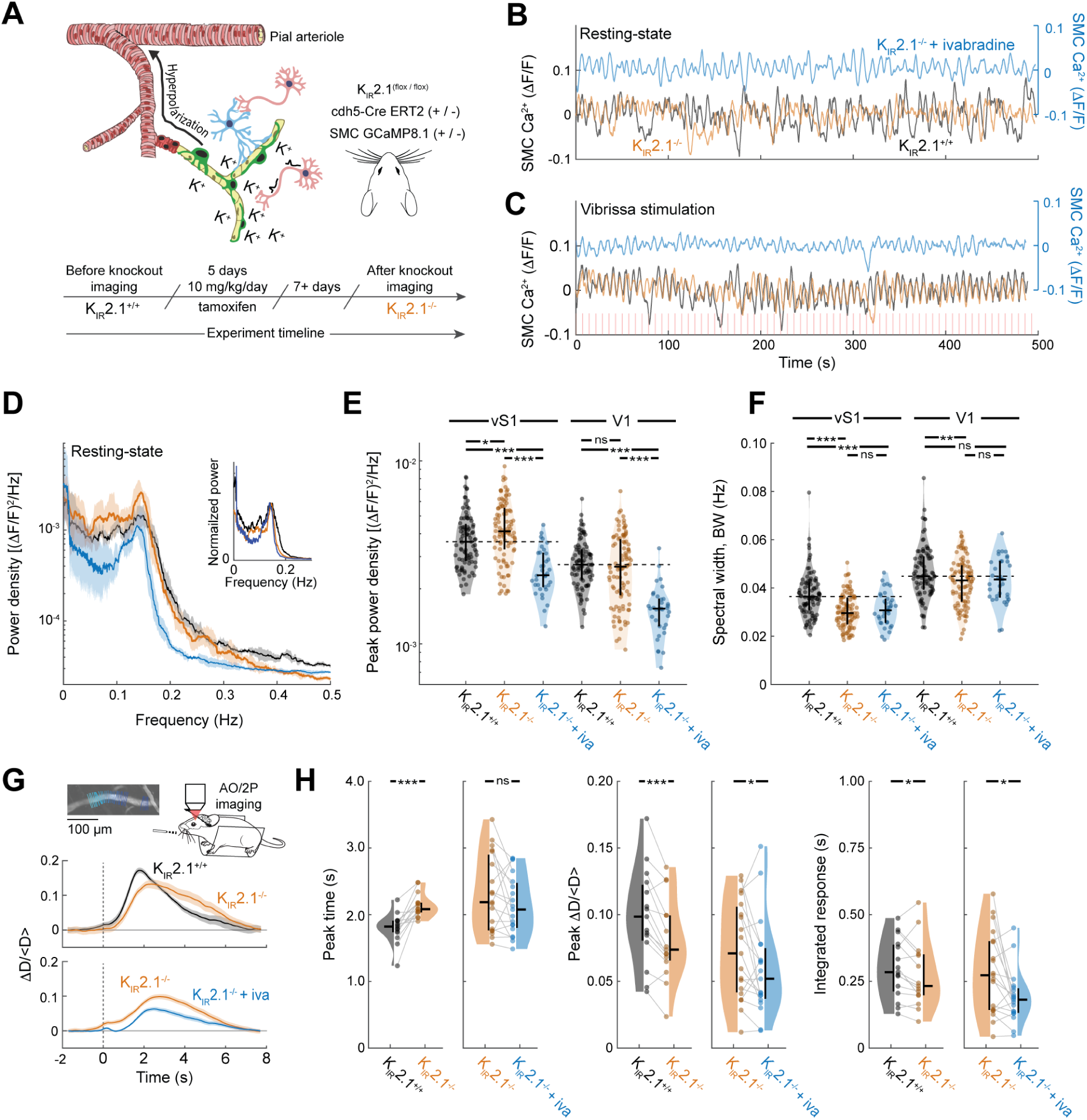
Alteration of pial arteriole activity during endothelial K_IR_2.1 knockout and HCN channel inhibition. **(A)** Schematic of a pial arteriole showing K_IR_2.1 channel-mediated hyperpolarization in endothelial cells (top; adapted from Sancho, M. and Nelson, M. T., 2021^104^). Conditional endothelial-specific K_IR_2.1 knockout mice (K_IR_2.1^flox/flox^; Cdh5-Cre ERT2^+/−^; SMC GCaMP8.1^+/-^) were generated by tamoxifen administration (10 mg/kg/day for 5 days), with wide-field calcium imaging performed before and ≥ 7 days after knockout induction. Where indicated, the HCN channel blocker ivabradine was co-administered. **(B)** Representative SMC Ca²⁺ (ΔF/F) traces (∼ 500 s) recorded from K_IR_2.1 WT (black), K_IR_2.1 KO (orange), and K_IR_2.1 KO + ivabradine (blue) animals during resting state. **(C)** Representative SMC [Ca²⁺] (ΔF/F) traces (∼ 500 s) recorded during 0.14 Hz vibrissa stimulation (red lines), plotted as in (B). **(D)** Power spectral density of resting-state SMC [Ca²⁺] signals across genotypes and treatment conditions (K_IR_2.1 WT, black; K_IR_2.1 KO, orange; K_IR_2.1 KO + ivabradine, blue). Each spectrum is from the same animal and are the mean ± 95% CI across four experiments. Inset shows the 0–0.3 Hz band on a linear scale, normalized to a common peak height, for comparison of peak widths between conditions. **(E)** Peak power density of resting-state vasomotor oscillations for arterioles in vS1 and V1 cortices across genotypes and treatment conditions. Each data point is one ∼ 500 s experiment. Bars indicate median, 25^th^, and 75^th^ percentiles. Asterisks denote * p < 0.05, ** p < 0.01, and *** p < 0.001 in all plots. **(F)** Resting-state vasomotor oscillation spectral width (Hz) for arterioles in vS1 and V1 cortices across genotypes and treatment conditions. Each data point is one ∼ 500 s experiment. Plotted as in (E). **(G)** Adaptive-optics two-photon (AO/2P) imaging of vasomotor dynamics during 0.10 Hz vibrissa stimulation. Top right: schematic of experiment. Top left: representative pial arterial vessel including lines to extract diameter via the full width at half maximum signal. Vibrissa 0.1 Hz stimulus-triggered average response (mean ± SEM) of pial arteriole diameter across conditions for two example experiments. **(H)** Summary of the stimulus-averaged pial diameter response during 0.1 Hz vibrissa stimulation. Bars indicate median, 25^th^, and 75^th^ percentiles.

Resting state vaso-oscillations remained after EC K_IR_2.1 knockout and after simultaneous EC K_IR_2.1 knockout and HCN channel blockade (**Figure 2B**). In each case, arteriole entrainment to periodic vibrissa stimulation remained, while the oscillation amplitude was reduced during simultaneous EC K_IR_2.1 knockout and HCN blockade (**Figure 2C**). After EC K_IR_2.1 knockout, resting state ∼ 0.1 Hz oscillations became more pronounced and remained during ivabradine administration, although with reduced spectral power (**Figure 2D**). Across experiments (188 experiments from 4 mice), the spectral power at the vasomotor peak increased in vS1 cortex but did not change significantly in V1 cortex after EC K_IR_2.1 knockout (**Figure 2E**; p = 0.01 and 0.20, one and two-tailed t-tests, respectively). Ivabradine administration reduced the peak power by 33 % and 41 %, in vS1 and V1 respectively, compared to before knockout (p = 5 x 10^-9^ and 9 x 10^-18^, one-tailed t-tests). In control mice, vasomotor power was unchanged after tamoxifen treatment in both vS1 and V1 cortices (**Figure S2A**; 66 experiments from 2 mice, p = 0.39 and p = 0.23, two-tailed t-tests, respectively). Post-knockout oscillations were spectrally sharper; across all experiments, the spectral width of resting-state vasomotion decreased from before to after EC K_IR_2.1 knockout in both vS1 and V1 cortices (**Figure 2F**; p = 1.4×10^-8^ and 0.004, respectively, one-tailed t-tests) whereas there was no significant change in spectral width in control animals after tamoxifen treatment (**Figure S2B**; p = 0.17 and 0.25, two-tailed t-tests, respectively) or after ivabradine administration (**Figure 2F**). We conclude that vasomotion near 0.1 Hz is generated intrinsically within the arteriole network. Endothelial K_IR_2.1 signaling does not produce this rhythm but broadens it, so removing that input yields a stronger, spectrally sharper oscillation while leaving entrainment to sensory drive intact. In contrast, HCN channels contribute to oscillation amplitude without contributing to its bandwidth.

### Endothelial K_IR_2.1 and HCN are necessary for neurovascular coupling

To characterize the functional contribution of EC K_IR_2.1 and HCN channels, we used two-photon microscopy to image the same pial arterioles before and after EC K_IR_2.1 knockout, with and without ivabradine. After EC K_IR_2.1 knockout pial arterioles displayed a blunted and delayed response to brief vibrissa stimulation, which was further diminished after ivabradine administration (**Figure 2G**). Across experiments (15 arterioles from 3 mice) the response time to peak increased after EC K_IR_2.1 knockout by 0.26 [0.15 – 0.50] s (**Figure 2H**; p = 3.1×10^-5^; median, 25^th^, and 75^th^ percentiles) whereas no difference in response time was observed in tamoxifen treated control animals (**Figure S2C**; median pairwise increase of -0.02 [-0.14 – 0.06] s, p = 0.39). After EC K_IR_2.1 knockout, the median decrease in normalized diameter response, Δ*D*⁄〈*D*〉, was 19 [12 – 27] % (p = 5.8×10^-4^) and the median decrease in integrated response was 14 [-2 – 19] % (**Figure 2H**; p = 0.04). This decrease in response amplitude was smaller than measured previously, in anesthetized mice, using one minute of continuous stimulation^34^.

In a separate cohort (19 arterioles from 4 mice), ivabradine administration decreased the normalized diameter response by 20 [-9 – 38] % and decreased the integrated response by 21 [-11 – 51] %, compared to EC K_IR_2.1 knockout alone (**Figure 2H**; p = 0.04 and 0.01 respectively). The change in peak time from before to after ivabradine administration was not significant (p = 0.10, two-tail Wilcoxon signed rank test; median change of -0.12 [-0.47 – 0.07] s). After EC K_IR_2.1 knockout the average arteriole diameter increased during both rest and sensory stimuli periods (**Figure S2D**) whereas in control animals (10 arterioles from 2 mice), tamoxifen treatment caused no change in average arteriole diameter (**Figure S2E**; p = 1 and p = 0.43 respectively, two-tail Wilcoxon signed rank tests). Control mice also exhibited no difference integrated response values after tamoxifen treatment, and an increase in normalized response amplitude (**Figure S2F**; p = 0.25 and 0.04, respectively). In total, knockout of EC K_IR_2.1 alone delayed and marginally reduced the response amplitude, while additional blockade of HCN channels was required to observe the lowest previously achievable response amplitudes^27^.

### Resting-state power is redistributed during vibrissa, visual, and RVLM stimulation

The intrinsic vaso-oscillation has a broad spectral peak (**Figures 1H** and **2F**), which opens the possibility of nonlinear interactions among the underlying arteriole oscillators as a function of stimulation strength and frequency relative to that of the underlying resting-state vasomotion (**Figure 1G** and **S1A**). To account for variability in resting-state vasomotion (**Figure 1D,E**), each trial consists of a resting-state measurement (250 to 500 s) followed immediately by periodic sensory stimulation (500 s). Stimulation near the vasomotor frequency entrains pial arterioles and produces characteristic peaks in the arteriole power spectrum (**Figure 3A**). To quantify the amplitude and timing of this response, we determine statistically if a given frequency component in the power spectral density can be replaced by delta functional^49,50^, i.e., a spectral "line" component. For the typical case of spectrally sharp responses solely at the drive frequency and harmonics, this is equivalent to fitting the temporal response with

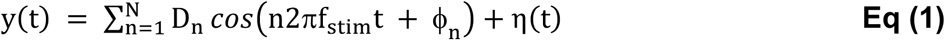

**Figure 3:**
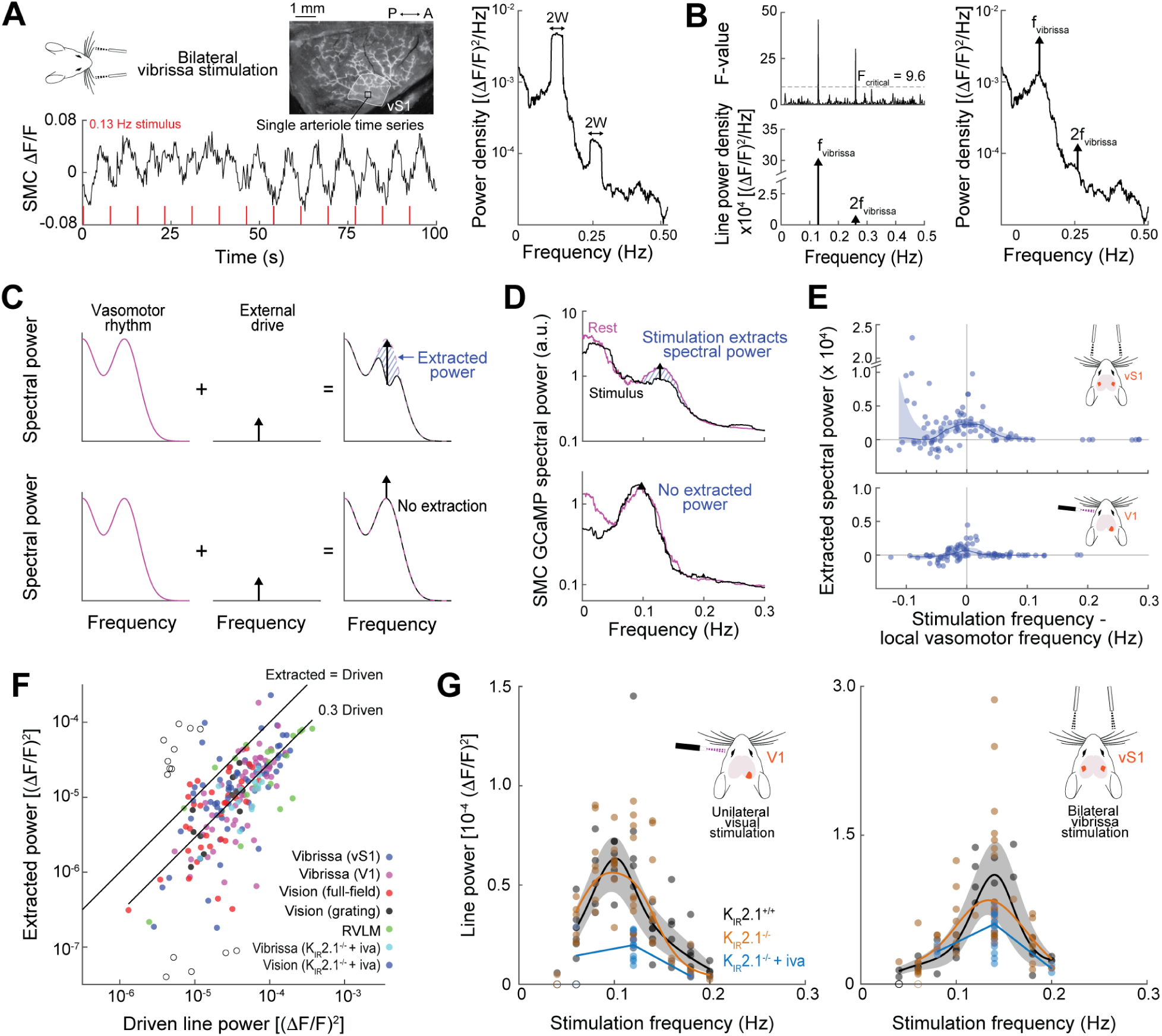
Spectral analysis and nonlinear properties of the stimulated pial response. **(A)** Example of vasomotor entrainment to bilateral vibrissa stimulation at 0.13 Hz. Left: wide-field fluorescence image of a pial arteriole in vS1 cortex (white outline) with the corresponding SMC GCaMP ΔF/F time series below, showing clear oscillations phase-locked to the 0.13 Hz stimulus (red tick marks). Right: power spectral density (PSD) of the SMC Ca²⁺ signal, exhibiting sharp peaks at the fundamental stimulation frequency (0.13 Hz) and its second harmonic (0.26 Hz), 2W = twice the Slepian taper half-bandwidth. **(B)** Frequency-domain analysis of sensory-evoked vasomotor responses. Top left: F-value spectrum from the line-component analysis; the dashed line indicates the F-value (F_critical_ = 9.6), above which spectral lines at f_vibrissa_ and 2f_vibrissa_ are statistically significant. Bottom left: line-power density spectrum showing discretely significant power values at f_vibrissa_ and 2f_vibrissa_. Right: PSD of the SMC GCaMP signal with peaks at f_vibrissa_ and 2f_vibrissa_ replaced by delta functionals. **(C)** Conceptual framework for spectral power extraction in non-linear and linear oscillatory systems. Top: non-linear system in which stimulation extracts broadband spectral power from nearby frequencies. The difference in broadband power from before to during stimulation, i.e. the extracted power, is highlighted in blue. Bottom: linear oscillatory system in which no power extraction occurs. **(D)** Demonstration of spectral power extraction in two example experiments. Top: SMC GCaMP PSD during rest (pink) and during periodic vibrissa stimulation (black), showing a reduction in broadband power flanking the stimulation frequency. Bottom: SMC GCaMP PSD during rest (pink) and periodic visual stimulation (black), showing negligible extracted power. **(E)** Extracted spectral power as a function of the difference between stimulation frequency and the local vasomotor frequency, for arterioles in vS1 cortex during vibrissa stimulation (top) and V1 cortex during visual stimulation (bottom). Each dot is one experiment; blue curves indicate the kernel-smoothed median and 95% confidence interval of the median. **(F**) Cross-modality and cross-region summary of extracted versus driven spectral power, pooling data from vibrissa stimulation (vS1, blue; V1, pink), full-field visual stimulation (red), visual grating stimulation (black), RVLM activation (green), and vibrissa and visual stimulation during K_IR_2.1 knockout + ivabradine. Open circle trials are excluded from the analysis. Reference lines indicate the loci where extracted power equals driven power (upper diagonal) and where extracted power equals 30% of driven power (lower diagonal). **(G)** Line-power as a function of stimulation frequency for unilateral visual stimulation (left) and bilateral vibrissa stimulation (right), with individual experiments (dots) and median ± 95% CI curves shown (K_IR_2.1 WT, black; K_IR_2.1 KO, orange; K_IR_2.1 KO + ivabradine, blue).

where the amplitude and relative phase of each harmonic line are given by *D_n_* and ϕ_n_, respectively, N is the highest harmonic, and η(t) is background activity, including vasomotion. The spectral power of the corresponding delta functional, referred to as the "line-power", is 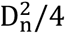 (**Figure 3B, STAR Methods**).

To assess how pial arterioles form the stimulated response, we exploit the concept of extracted spectral power, i.e., the difference in background spectral power near the stimulus frequency between stimulated and resting conditions^51,52^ (**Figure 3C**). For a system of non-linear coupled oscillators, stimulation at one frequency can extract power from oscillators at neighboring frequencies (top, **Figure 3C,D**) so that the baseline spectrum is diminished. In contrast, for a linear system the stimulated response simply adds to an unaltered baseline response (bottom, **Figure 3C,D**). We consider extracted power as a metric for nonlinearity. During both vibrissa and visual stimulation, arterioles in the associated primary sensory cortices extract significant energy at their local vasomotor frequency (**Figure 3E**; p < 1×10^-4^ and p = 1×10^-3^ for vS1 and V1 cortices, respectively, one-tail bootstrap tests). The extent of extraction is a function of response amplitude at the drive frequency. During epochs of vibrissa stimulation that lead to line-power larger than ∼ 10^-5^ (ΔF/F)^2^, extracted and line-power become significantly correlated (**Figure S3A**). This correlation is also apparent during visual stimulation with either flashing illumination (**Figure S3B**) or with a drifting grating (**Figure S3C,D**), and via activation of RVLM (**Figure S3E**). A summary across all data sets, including experiments during simultaneous EC K_IR_2.1 knockout and HCN blockade, shows a correlation of extracted energy with driven energy that has an essentially linear slope of 0.93 ± 0.07, with extracted energy accounting for 0.30 ± 0.02-times the driven energy (**Figure 3F**; mean ± SE). The observation that a fixed fraction of power is extracted over a wide range of conditions and cortical locations implies energy extraction is a universal aspect of non-linear vasodynamics.

Resting-state variability partially determines the stimulus-response amplitudes. Thus, without knowledge of the resting-state activity, large-amplitude stimulated responses may be erroneously attributed to stronger neural signaling alone. By example, we consider the stimulated arteriole response during EC K_IR_2.1 knockout, which is associated with increased resting-state power compared to before knockout (**Figure 2D,E**). Across stimulation frequencies, the periodic component of the response is unchanged from before to after EC K_IR_2.1 knockout for both vibrissa and visual stimulation, presumably reflecting extraction of upregulated resting-state power that compensates for decreased neurovascular signaling (**Figures 3G** and **2D,E**). Ivabradine administration decreases the median periodic response by 60 % and 45 %, when tested at 0.12 Hz and 0.14 Hz for visual and vibrissa stimulation respectively. This change presumably reflects resting-state power that is reduced during this condition and is unable to overcome this extreme neurovascular perturbation (**Figures 3G** and **2D,E,G**).

### Pial arterioles resonantly enhance underlying neural activity near the vasomotor frequency

Resonance is a universal characteristic of forced systems with an intrinsic time scale. Thus, given the observation that the pial arteriole network undergoes intrinsic oscillations *in vivo*, we hypothesize that resonance phenomena shape the pial arteriole response to neural activity. We first consider amplitude enhancement for the case of visual stimulation at frequencies in the vicinity of the vasomotor frequency. Periodic visual stimulation elicits a periodic neural response that is centered within the area of V1 cortex, as shown in the example of **Figure 4A**, as well as drives periodic arteriole activity in the pial arterioles overlying V1 cortex, as shown for the same example experiment of **Figure 4B**. The amplitude of periodic neural activity varies with stimulation frequency with a broad maximum at 0.13 Hz (88 experiments in 12 mice) (**Figure 4C**). In the same experiments, the arteriole response sharply peaks at 0.11 Hz, consistent with a resonance at the network’s peak resting-state frequency (**Figures 1G** and **4D**). To determine if this arteriole response is caused by arterial resonance or an ultra-low frequency variation in the neural response, we consider each system’s response ratio from on- to off-resonance. From 0.04 Hz to peak, neural power increases by 2.8-times, whereas arteriole power increases by a factor of 3.9 (**Table S1; Figure 4C,D**). When normalized on an experiment-by-experiment basis by neural power, arteriole power peaks with an enhancement factor of 1.9 from 0.04 Hz to 0.10 Hz (**Table S1; Figure S4A**). We conclude that the pial arterioles show a strong enhancement of the underlying neural activity near a vascular resonance at the vasomotor frequency.

**Figure 4:**
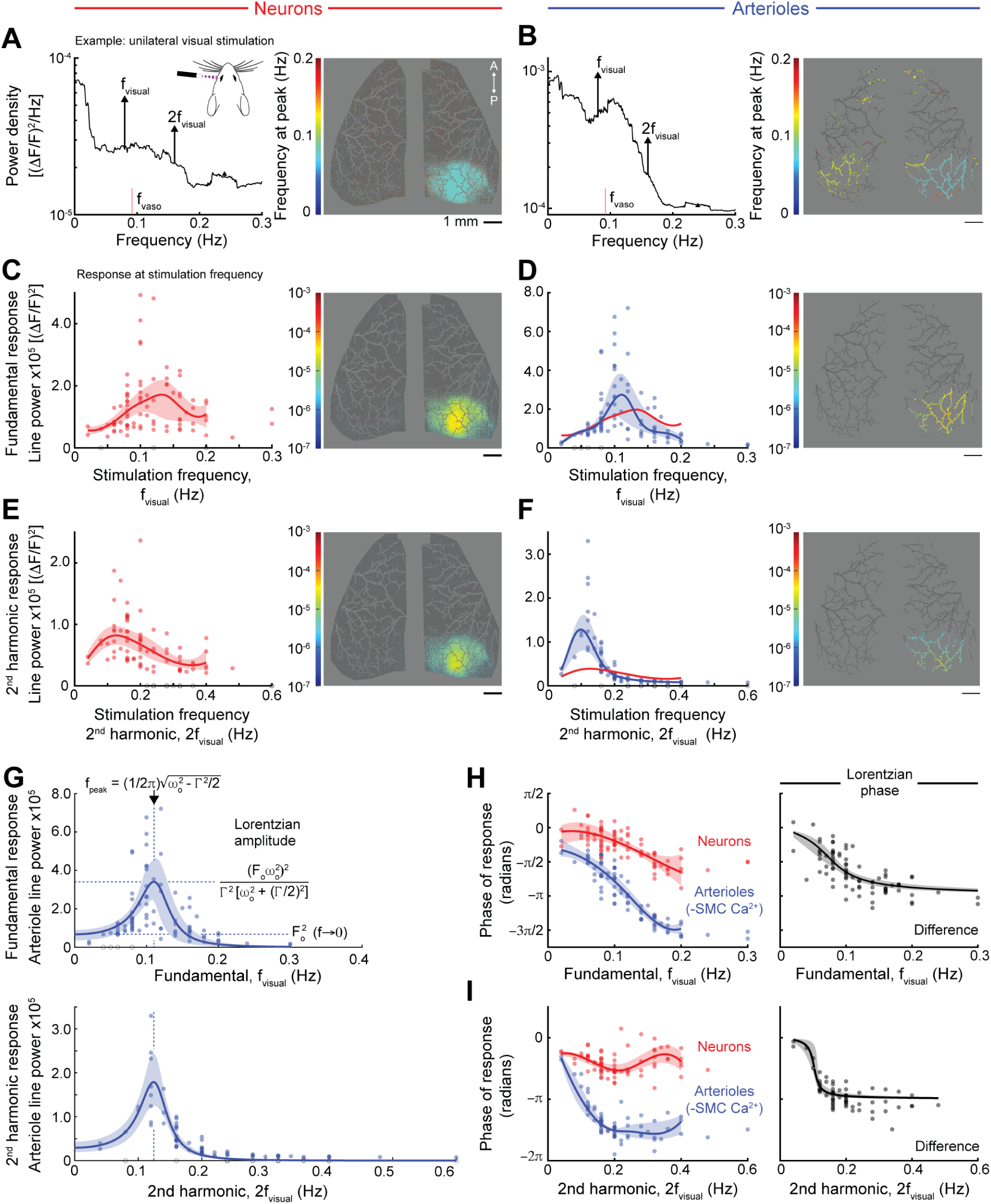
Pial arterioles amplify visually evoked neural activity near the resting-state vasomotor frequency. **(A)** Left: representative power spectral density (PSD) of jRGECO neuronal Ca²⁺ signal during unilateral visual stimulation at f_visual_ = 0.08 Hz. Spectra are averaged over pixels in V1 cortex. The local vasomotor frequency (f_vaso_) is indicated by a red tick. Inset schematic shows the unilateral visual stimulation setup. Right: Dorsal cortical map of frequency at maximum line-power. Neuronal pixel color-coded dark gray denote regions in the mask whose signal did not contain any significant line components. Line component responses cluster in the contralateral posterior (visual) cortex at stimulation frequencies (color code: cyan – 0.08 Hz; dark blue – 0; red – 0.2 Hz). **(B)** Left: Representative arteriole SMC GCaMP PSD during the example experiment shown in (A). Right: cortical vascular map with arterioles color-coded by their frequency at maximum line-power. Arterioles with no significant line components are colored dark gray. **(C)** Neural line-power as a function of the stimulus fundamental frequency. Left: Average V1 cortical line-power across all visual stimulus experiments with kernel-smoothed median and 95 % CI for the median shown (red dots: individual experiments). Open circles denote experiments that did not exhibit more significant line components at the stimulus frequency than expected by chance (**STAR Methods**). Right: Spatial map of neuronal line-power at 0.08 Hz for the example experiment shown in (A). **(D)** Arteriole line-power as a function of the stimulus fundamental frequency. Left: Average V1 cortical line-power for all visual stimulus experiments, plotted as in (C). Each blue dot denotes an individual experiment. The neural median curve (red) is overlaid after scaling by the ratio of arteriole to neural power at 0.04 Hz, for comparison (**STAR Methods**). Right: Spatial map of arteriole line-power at 0.08 Hz for the example experiment shown in (A). **(E)** Neural 2^nd^ harmonic line-power as a function of the stimulus frequency 2^nd^ harmonic. Left: Average V1 line-power at 2^nd^ harmonic frequencies across all visual stimulus experiments, plotted as in (C). Right: Spatial map of neuronal line-power at 0.16 Hz for the example experiment shown in (A). **(F)** Arteriole 2^nd^ harmonic line-power as a function of the stimulus frequency 2^nd^ harmonic. Left: Average V1 line-power at 2^nd^ harmonic frequencies across all visual stimulus experiments, plotted as in (D). The neural median curve (red) is overlaid after scaling by the ratio of arteriole to neural power at 0.18 Hz, for comparison (**STAR Methods**). Right: Spatial map of arteriole line-power at 0.16 Hz for the example experiment shown in (A). **(G)** Lorentzian fit, ± 95 % CI, to the distribution of arteriole fundamental (top) and 2^nd^ harmonic (bottom) responses as a function of drive frequency and drive frequency 2^nd^ harmonic. The peak frequency is given by f_peak_ = (1/2π)√(ω₀² − Γ²/2), indicated by the dashed vertical line. Horizontal dashed lines mark the Lorentzian amplitude 1/(Γ√(ω₀² − (Γ/2)²) at the peak and the low-frequency asymptote (1/ω₀² as f → 0), providing quantitative estimates of the oscillator’s resonant gain and damping. (**Eq 5**). **(H)** Neural and arteriole line component response phase, averaged over pixels in V1 cortex. Left: phase of response (radians) as a function of fundamental stimulation frequency (Hz) for neurons (red) and arterioles (−SMC [Ca²⁺], blue), with kernel-smoothed median and 95 % CI. Right: the neuron–arteriole phase difference fit to the equation for a linear oscillator (black curve ± 95 % CI, **Eq 6**) as a function of fundamental frequency, quantifying the phase lag introduced by neurovascular coupling. **(I)** Neural and arteriole line component response phase at the stimulation frequency 2^nd^ harmonic, plotted as in (H).

Flickering full field visual stimuli may only activate limited neural populations in comparison to naturalistic stimuli^53^. To address this concern, we performed visual stimulation experiments with a moving square wave grating (**Figure S3C**). The median response to a grating was larger than responses to full-field stimulation in terms of both arteriole and neural power (**Figure S4B**). The largest-amplitude responses had the largest extracted power values and conformed well to the extracted power versus line-power relationship for full field visual stimuli (**Figure S3D**). We conclude that full-field stimuli can be used to probe the low-amplitude regime, while moving gratings can be used to drive larger arteriole responses in the visual system.

Periodic stimuli elicit neural activity at various harmonics through either harmonics present in the stimulus itself or nonlinearities in neural processing^54^. Additional vascular nonlinearities likely contribute to harmonics in the arteriole response^55,56^. As expected, our non-sinusoidal periodic stimulation paradigm elicits periodic neural activity at the harmonics of the stimulation frequency (**Figure 4A,E**) that entrain the overlying pial arterioles (**Figure 4B,F**). The 2^nd^ harmonic response peaks at 0.13 Hz and 0.10 Hz for neurons and arterioles, respectively (**Table S1**), and is enhanced by 1.1- and 3.7-times from 0.18 Hz to the frequency of maximum response, for neurons and arterioles respectively (**Table S1**), mirroring the system’s response at the fundamental frequency (**Figure 4G**). A 3.2-fold enhanced 2^nd^ harmonic arteriole response is seen clearly in the normalized response (**Table S1; Figure S4C**).

Both the fundamental and 2^nd^ harmonic arteriole amplitude responses to visual stimulation are likely to be only weakly non-linear, as judged from the relatively low level of extracted power (**Figure 3F**). As such, it is useful to observe if they can be described by the equation for a linear oscillator^48^ to good approximation (**Figure 4G; Eqs 4-5**). In this model, the temporal delay between the oscillator and external forcing follows an arctangent function (**Eq 6**); near zero frequency the oscillator tracks the forcing without delay, on resonance the oscillator lags by a phase of *π*⁄2 radians (a time delay of 1/4f_peak_) and at high frequencies the response becomes anti-phasic with the forcing. Accordingly, we consider the phase delay from neural activity to arteriole dilation using the average line component phase in V1 cortex (**Eq 1**). The phase of neural activity decreases roughly linearly with frequency, commensurate with our 2-s duration stimulation, and the observed arteriole phase decreases linearly before plateauing at frequencies above 0.2 Hz (**Figure 4H**). The difference between the two-phase curves yields the phase delay from neural activation to arteriole dilation, which plateaus at a phase of *π* radians and is well fit by the arctangent function (**Eqs 4-6, STAR Methods**). This relationship also holds for activity at the 2^nd^ harmonic (**Figure 4I**). The phase of variations in arteriole diameter, inferred using the previously characterized phase relationship between SMC [Ca^2+^] and diameter^46^, follows the SMC [Ca^2+^] trend with an additional delay (**Figure S4D,E**).

We next tested if resonance is a common feature of the pial arterioles by probing the network’s response to vibrissa sensory stimulation, which typically has a higher level of driven and extracted power (**Figure 3F**). Periodic vibrissae stimulation via air puffs elicits a periodic neural response that extends beyond the boundaries of primary vibrissa cortex (**Figure 5A**). Qualitatively, the peak arteriole response follows the stimulus when analyzed in terms of phase-locked cortical area (**Figure 5B**). When repeated for many frequencies near 0.1 Hz (95 experiments and 16 mice), neural activity power in vS1 cortex displays a moderate trend as a function of stimulation frequency (**Figure 5C**). Arteriole power peaks at 0.13 Hz and is enhanced by 5.3-times from off- to on-resonance (**Table S1**; **Figure 5D**). Neural power increased by 2.2-times across the same range, indicating that the observed arteriole resonance is not caused by a resonance in the neural sensory response. When normalized by neural power, the arteriole response peaks at 0.13 Hz, with a 2.3-times enhancement from 0.04 Hz to 0.13 Hz, further supporting an ∼ 0.1 Hz arteriole resonance (**Figure S5A**).

**Figure 5:**
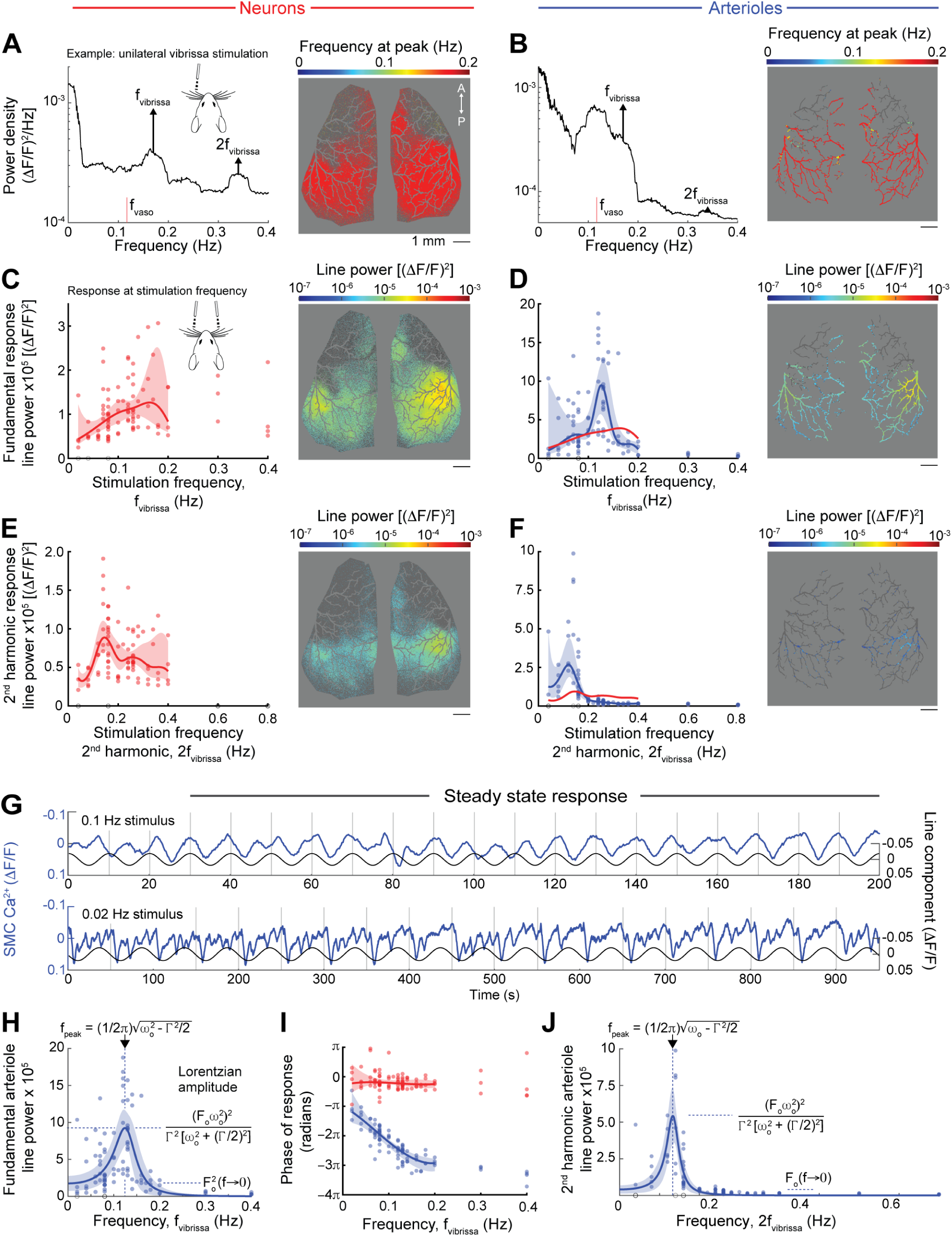
Pial arterioles resonantly amplify vibrissa neural activity during vibrissa stimulation. **(A)** Left: representative power spectral density (PSD) of jRGECO neuronal Ca²⁺ signal during unilateral vibrissa stimulation at f_visual_ = 0.17 Hz. Spectra are averaged over pixels in vS1 cortex. The local vasomotor frequency (f_vaso_) is indicated by a red tick. Inset schematic depicts the unilateral stimulation scheme. Right: Dorsal cortical peak frequency map with each neuronal pixel color-coded by its frequency at maximum line-power (dark blue – 0 Hz; dark red – 0.2 Hz). **(B)** Left: representative arterial SMC GCaMP PSD during the unilateral vibrissa stimulation experiment show in (A). Right: dorsal cortical maximum vascular map with arterioles color-coded by their frequency at maximum line-power (dark blue – 0 Hz; dark red – 0.2 Hz). **(C)** Neural line-power as a function of the stimulus fundamental frequency. Left: Fundamental response line-power for individual neuronal recordings (red dots) as a function of stimulation frequency (Hz), with smoothed median and 95% CI (red curve and shaded band). Right: Spatial map of neuronal line-power at 0.17 Hz for the example experiment shown in (A). **(D)** Arteriole line-power as a function of the stimulus fundamental frequency. Left: Average vS1 cortical line-power for all vibrissa stimulus experiments, plotted as in (C). Each blue dot denotes an individual experiment. The neural median curve (red) is overlaid after scaling by the ratio of arteriole to neural power at 0.04 Hz, for comparison (**STAR Methods**). Right: Spatial map of arteriole line-power at 0.17 Hz for the example experiment shown in (A). **(E)** Neuron 2nd harmonic line-power as a function of the stimulus frequency 2^nd^ harmonic. Left: Average vS1 line-power at stimulus 2^nd^ harmonic frequencies for all visual stimulus experiments, plotted as in (C). Right: Spatial map of neuronal line-power at 0.34 Hz for the example experiment shown in (A). **(F)** Arteriole 2nd harmonic line-power as a function of the stimulus frequency 2^nd^ harmonic. Left: Average vS1 line-power at 2^nd^ harmonic frequencies across all visual stimulus experiments, plotted as in (D). The neural median curve (red) is overlaid after scaling by the ratio of arteriole to neural power at 0.18 Hz, for comparison (**STAR Methods**). Right: Spatial map of arteriole line-power at 0.34 Hz for the example experiment shown in (A). **(G)** Entrained arteriole Ca²⁺ responses at two stimulation frequencies. Time traces of SMC [Ca²⁺] (blue, left y-axis) and the line component (right y-axis) are shown for two representative stimulation conditions. Gray vertical ticks denote stimulus time points. **(H)** Lorentzian fit, ± 95 % CI, to the distribution of arteriole fundamental responses to vibrissa stimulation as a function of drive frequency. **(I)** Neural and arteriole line component response phase, averaged over pixels in vS1 cortex. **(J)** Lorentzian fit, ± 95 % CI, to the distribution of arteriole 2^nd^ harmonic responses to vibrissa stimulation as a function of drive frequency 2^nd^ harmonic.

As in the case of visual stimuli, our periodic vibrissa stimulation paradigm elicits periodic neural activity at the harmonics of the stimulation frequency (**Figure 5A,E**) that entrain the overlying pial arterioles (**Figure 5B,F**). The 2^nd^ harmonic response peaks at 0.14 Hz and 0.12 Hz for neurons and arterioles, respectively, and is enhanced by 1.2 and 3.5 times from 0.18 Hz to the frequency of maximum response, for neurons and arterioles respectively, mirroring the system’s response at the fundamental frequency (**Table S1**). The normalized 2^nd^ harmonic arteriole response peaks at 0.07 Hz and is enhanced by 4.9-times from 0.18 Hz to 0.07 Hz (**Figure S5B**). All told across visual and vibrissa cortices, the neural representation of sensory stimulation is demonstrably enhanced by the vasomotor representation for stimuli presented near the nominally 0.05 to 0.2 Hz range of resting state vasomotion.

The response of the vibrissa system to stimulation near or above ∼ 0.07 Hz followed that expected for a resonator (**Figure 5G**). Yet for stimuli near or below ∼ 0.07 Hz the response in select trials was complex; each cycle contained a strong constriction followed by ∼ 0.1 Hz ringing (**Figure 5G**). The novel constriction is of unknown origin, while the ringing appears as the natural response to a perturbative "kick" to the vaso-oscillator. This leads to a high amplitude response at low frequencies compared to those seen in V1 cortex (**Figure 4G**) as is expected for a linear oscillator (cf **Figures 4G** and **5H**). More so, the calculation of the phase shift is distorted by the large constriction, such that the calculated arteriole phase lag spans the entire 2*π* range (**Figure 5I**). A similar argument holds for the 2nd harmonic of the response (**Figures 5J** and **S5C**).

### Stimulation at the resting-state frequency preferentially entrains pial arteriole territory

Stimulus-evoked entrainment of a cortical area is a widely used metric of functional boundaries and connectivity in fMRI^2^. So far, we have analyzed activity within the stimulated primary cortical areas, however resonant enhancement may also increase the likelihood of entraining neighboring arterioles. For this reason, we hypothesize that the spatial extent of entrainment depends on the spectral content of the stimuli relative to the resting-state vasomotor frequency. To assess areal entrainment, we measured the spatial extent of periodic activity across the cortical mantle, using line-power at the stimulus frequency, or at the second harmonic of this frequency, as a metric (**Eq 1; STAR Methods**). Indeed, close to the resting-state vasomotor frequency, vibrissa stimulation is found to entrain pial arterioles well beyond the boundary of neural activation, with stimulation at the resting-state frequency entraining the entire cortical mantle (**Figure 6A**). This aspect of spatial saturation was not observed during visual stimulation, which is generally weaker in strength (**Figure 3F**), in which case pial arterioles entrained to neural stimulation only slightly enlarged their areal representation (**Figure 6B**).

**Figure 6:**
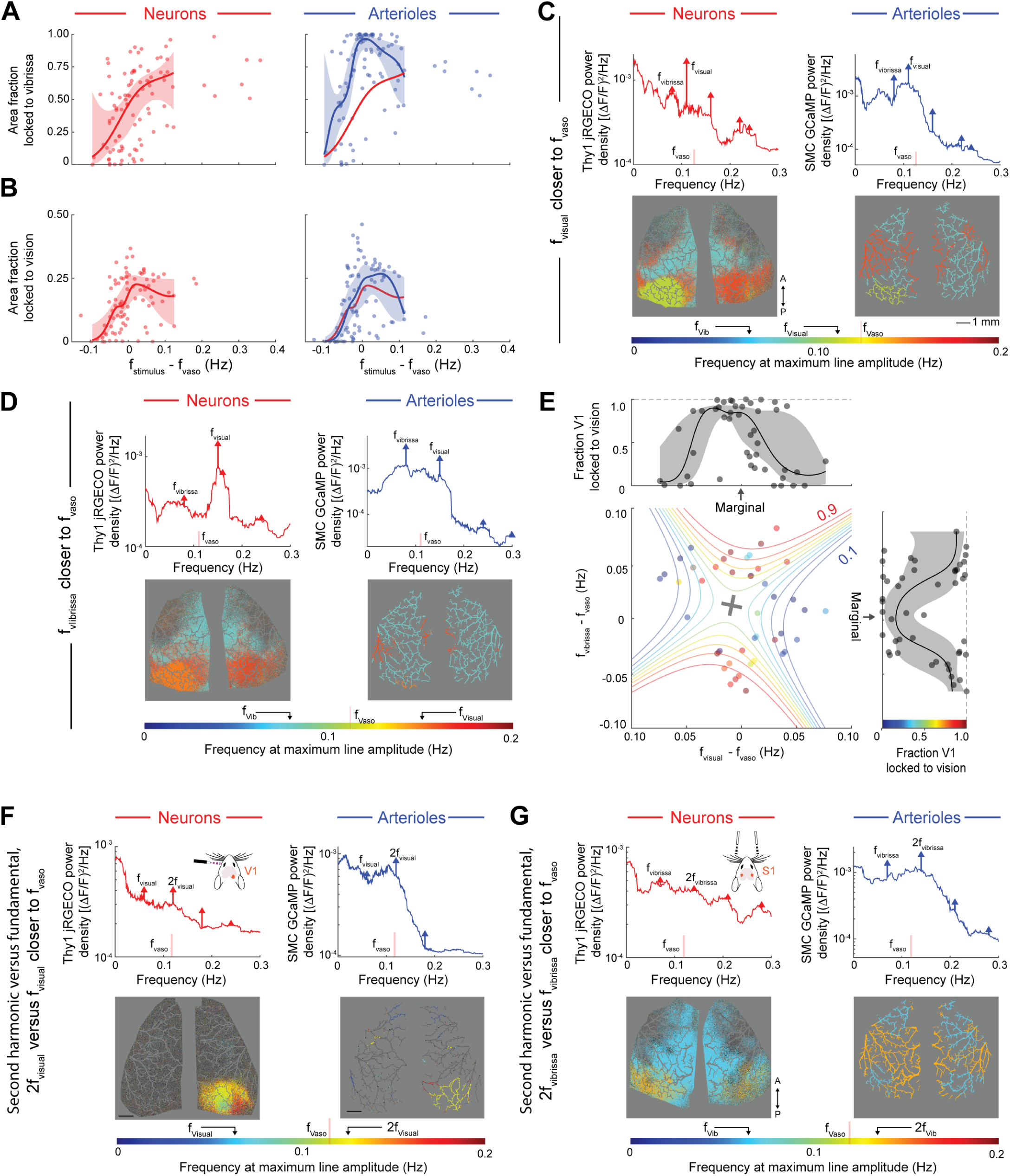
Pial responses to visual and vibrissa stimulation compete to entrain cortical territory. **(A)** Fraction of the contralateral hemisphere (or fraction of both hemispheres in the case of bilateral stimulation) entrained to uni/bilateral periodic vibrissa stimulation for the pial arterioles and neurons. Left: neuronal area fraction locked to f_vibrissa_ versus the frequency proximity to the resting-state vasomotor frequency (f_vibrissa_ − f_vaso_ (Hz)). Right: arteriole area fraction locked to stimulation frequency versus the frequency proximity to the resting-state vasomotor frequency (f_vibrissa_ − f_vaso_ (Hz)). Dots represent individual experiments and curves denote the kernel-smoothed median ± 95% CI. **(B)** Fraction of the contralateral hemisphere entrained to unilateral periodic full-field visual stimulation for the pial arterioles and neurons, plotted as in (A). **(C)** Example competition experiment between visual and vibrissa stimulation, in which the visual stimulation frequency (f_visual_) is closer to vasomotion (f_vaso_) than the vibrissa stimulation frequency (f_vibrissa_). Here f_vaso_ = 0.13 Hz (yellow), f_visual_ = 0.11 Hz (green), and f_vibrissa_ = 0.08 Hz (cyan). Top row: SMC-GCaMP8.1 arteriole (right) and Thy1-jRGECO neuronal (left) PSD averaged over V1 cortex. Bottom row: frequency at maximum line-power maps. **(D)** Example inverted competition experiment between visual and vibrissa stimulation with f_vaso_ = 0.11 Hz (green), f_visual_ = 0.15 Hz (orange), and f_vibrissa_ = 0.08 Hz (cyan). Plotted as in (C). **(E)** Two-dimensional summary of arteriole spatial entrainment during simultaneous visual and vibrissa stimulation. Each dot denotes one experiment, colored by the fraction of arterioles in V1 cortex locked to f_visual_. Areal locking fraction is plotted versus the proximity of each stimulus to the vasomotor frequency (abscissa: f_visual_ – f_vaso_, ordinate: f_vibrissa_ − f_vaso_). Overlaid are iso-fraction contours from a quadratic surface fit. Marginal plots are the area locking fraction one-dimensional distributions, with kernel-smoothed median ± 95% CI. **(F)** Example unilateral full-field visual stimulation experiment in which the visual stimulation 2^nd^ harmonic frequency (2f_visual_) is closer to vasomotion (f_vaso_) than the visual stimulation fundamental frequency (f_visual_). Here f_vaso_ = 0.12 Hz and f_visual_ = 0.06 Hz. Top row: SMC-GCaMP8.1 arteriole (right) and Thy1-jRGECO neuronal (left) PSD, averaged over V1 cortex. Bottom row: frequency at maximum line-power maps. Scale bar, 1mm. **(G)** Example bilateral vibrissa stimulation experiment showing pial entrainment to the stimulus 2^nd^ harmonic frequency, despite predominant neural activity at the stimulation fundamental frequency. Here f_vaso_ = 0.12 Hz and f_vibrissa_ = 0.07 Hz. Top row: SMC-GCaMP8.1 arteriole (right) and Thy1-jRGECO neuronal (left) PSD, averaged over vS1 cortex. Bottom row: frequency at maximum line-power maps.

Neuronal input based on environmental stimuli has both a broad-spectrum and encompasses multiple sensory channels, e.g., visual and tactile. How does arteriole resonance influence the pial network’s response to diverse stimuli? Here we use simultaneous presentation of visual and vibrissa stimuli at different frequencies and probe the spatial extent of arteriole entrainment. When visual stimulation is presented closer to the vasomotor frequency than vibrissa stimulation, arterioles overlying V1 cortex lock completely to the visual stimulation frequency and are locked one-to-one with neural activity in V1 cortex (**Figure 6C**). However, when vibrissa stimulation is presented closer to the vasomotor frequency, arterioles in V1 cortex become locked to the vibrissa frequency despite dominant V1 cortical neural activity at the visual stimulation frequency (**Figure 6D**). In competition, we observed that on resonance visual stimulation reliably locks arterioles in V1 cortex, but when vibrissa stimulation is closer to vasomotion, activity at the vibrissa frequency overtakes vasoactivity in these arterioles (47 experiments from 13 mice) (**Figure 6E**).

As a second test of competition, we consider the case of the fundamental versus second harmonic activity in proximity to the vasomotor frequency. We found that arterioles preferentially lock to the stimulus second harmonic when it is closer to the vasomotor frequency than the stimulus fundamental (**Figure 6F**). Interestingly, for low frequency visual stimulus experiments (f_drive_ ∼ 0.05 Hz) the largest neural response often occurs at the stimulus 2^nd^ harmonic (**Figure 6F**). This is consistent with the resonance-like neural response near 0.1 Hz when analyzed in terms of cortical area (**Figure 6B**). Competition between harmonic components is general, as arterioles lock to the stimulus 2^nd^ harmonic during low frequency vibrissae stimulation as well (**Figure 6G**).

### RVLM competes with sensory input for pial arteriole territory

How general is the issue of neuronal control of cortical vasomotion and competition among different neural inputs for control of arteriole dynamics? We test the consequence of driving a vasoactive medullary nucleus, the RVLM^41^, on pial arteriole blood flow in the context of ongoing sensory cortical activity. These experiments make use of mice expressing SMC GCaMP8.1 and dβh-Cre. We use a Cre-dependent AAV to express the red-shifted opsin ChRmine^47^ in adrenergic cells in the RVLM (**Figure 7A**). We first performed a set of experiments to titrate the optogenetic power and classify animals as either responders or non-responders based on their functional response (**Figure S6A,B**). Post-mortem histology confirmed that non-responding animals lacked ChRmine expression in RVLM; these animals were excluded from further analysis (**Figure S6B**). When presented with simultaneous vibrissa sensory stimulation, arranged to be equidistant in frequency from the resting-state vasomotor frequency, the sensory stimulation dominates pial arteriole entrainment when RVLM activation is weak (**Figure 7B**, middle). However, as RVLM activation strength is increased, subcortical input eventually outcompetes sensory input for the entrainment of pial arterioles across the cortical mantle (**Figure 7B**, right).

**Figure 7:**
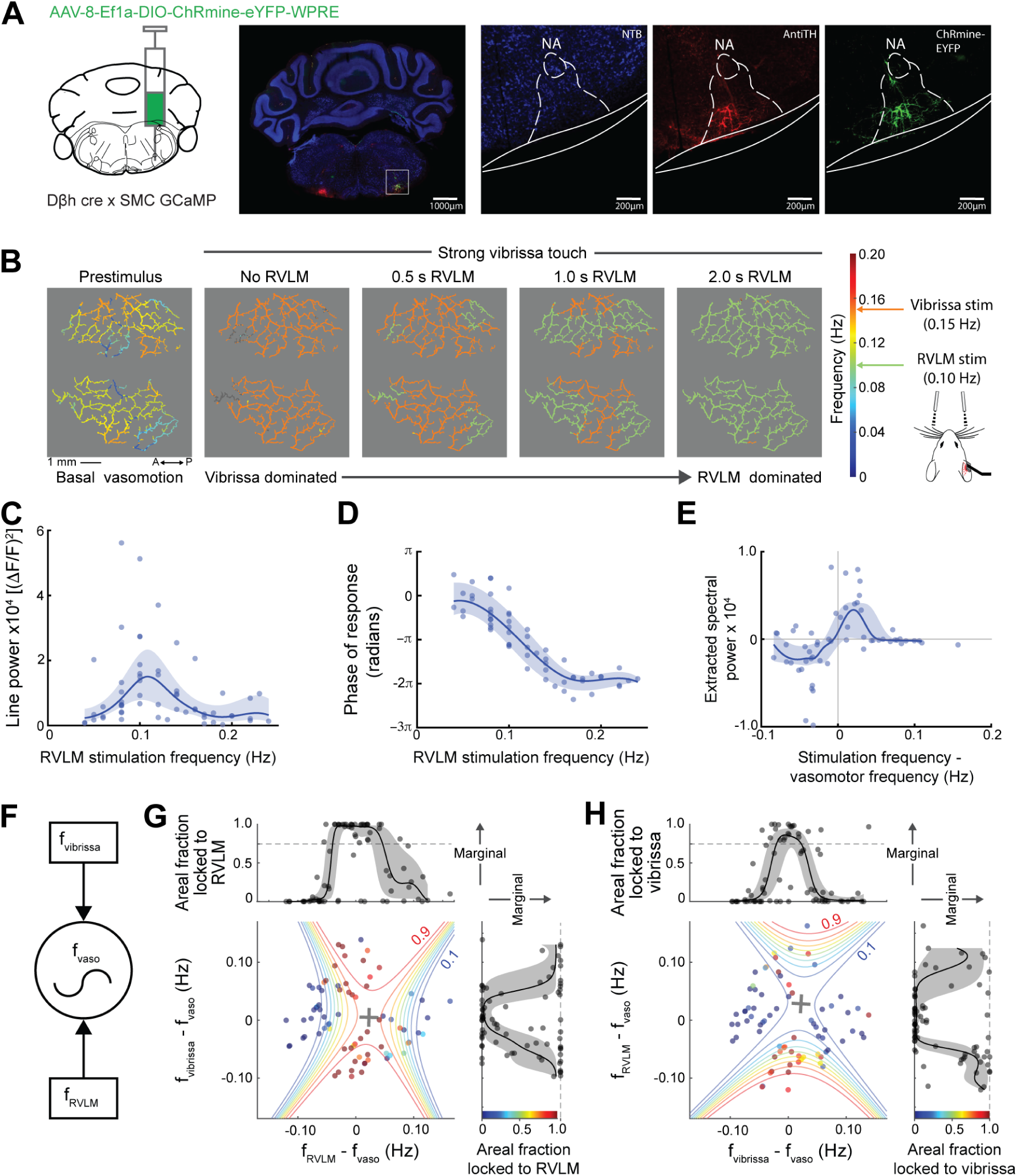
Optogenetic activation of adrenergic RVLM neurons entrains cortical arterioles and competes with vibrissa-driven entrainment in a frequency-dependent manner. **(A)** Viral strategy and histological validation of RVLM-specific ChRmine expression in Dβh-Cre × SMC-GCaMP mice. Left: schematic depicting stereotaxic injection of AAV-8-Ef1a-DIO-ChRmine-eYFP-WPRE into the rostral ventrolateral medulla (RVLM) of Dβh-Cre mice that co-express GCaMP in SMCs. Center left: coronal brain section with white box indicating the RVLM injection site co-stained with anti-tyrosine hydroxylase immunostaining (AntiTH, red), ChRmine-eYFP expression (green), and counter stain (NTB, Neuro trace blue). Remaining panels: individual channel images of the nucleus ambiguus (NA) and RVLM (dashed outline) injection site showing co-localization of tyrosine hydroxylase and ChRmine-eYFP. **(B)** Competition experiments between RVLM and vibrissa stimulation, with increasing RVLM stimulation durations. Left: pre-stimulus period with arterioles colored by their frequency at maximum total power. Remaining panels: simultaneous RVLM and vibrissa stimulation experiments, in which RVLM stimulation duration is increased from 0 s to 2 s. Here arterioles are colored by their frequency at maximum line-power. **(C)** Cortex-averaged arteriole line-power plotted against RVLM stimulation frequency for individual experiments (blue dots), with kernel-smoothed median ± 95% CI (blue curve and shaded band). Here, experiments consisted of RVLM stimulation only. **(D)** Arteriole response phase vs. RVLM stimulation frequency, averaged over all arterioles, for the same dataset presented in (C). **(E)** Extracted spectral power plotted as a function of the difference between RVLM stimulation frequency and vasomotor frequency (Hz). Blue dots are individual experiments, with the kernel-smoothed median ± 95% CI curve overlaid. **(F)** Diagram illustrating that both vibrissa sensory drive (f_vibrissa_) and RVLM cortical projection input (f_RVLM_) converge onto the vasomotor oscillator (f_vaso_), which gates cortical arteriole entrainment based on proximity to the oscillator’s natural frequency. **(G)** Two-dimensional summary of arteriole spatial entrainment during simultaneous vibrissa and RVLM stimulation. Each dot denotes one experiment, colored by the fraction of arterioles locked to f_RVLM_. Areal locking fraction is plotted versus the proximity of each stimulus to the vasomotor frequency (abscissa: f_RVLM_ – f_vaso_, ordinate: f_vibrissa_ − f_vaso_). Overlaid are iso-fraction contours from a quadratic surface fit. Marginal plots show area locking fraction one-dimensional distributions, with kernel-smoothed median ± 95% CI. One outlier point with zero areal fraction was excluded from the y-axis marginal fit (right panel). **(H)** Two-dimensional summary of arteriole spatial entrainment, for the same dataset plotted in (G), here with points colored by the fraction of arterioles locked to f_vibrissa_. One outlier point with an areal fraction of ∼ 1 was excluded from the x-axis marginal fit (top panel) and two outlier points at approximately f_RVLM_ - f_vaso_ = ± 0.12 Hz were excluded from the y-axis marginal fit (right panel).

To further examine competition for cortical territory by RVLM versus sensory stimulation, we performed RVLM activation experiments as a function of frequency relative to the resting-state vasomotor frequency. The pial arteriole response during RVLM activation is enhanced by 6.6-times from off- to on-resonance and peaks at 0.11 Hz (**Table S1**; **Figure 7C**). Notably, the median peak response on resonance was 1.6- and 5.4-times larger than those observed during vibrissa and full-field visual stimulation respectively (**Figure 7C**; peak values are 15 [7.7, 23] x 10^-5^ (ΔF/F)^2^, 9.5 [5.7, 12] x 10^-5^ (ΔF/F)^2^, and 2.8 [1.5, 3.8] x 10^-5^ (ΔF/F)^2^ with [95% CI] for RVLM, vibrissa, and visual stimulus respectively). The phase response during RVLM activation spanned the entire 2*π* range, in part due to constrictive peaks analogous to those observed during vibrissa stimulation (**Figures 7D** and **S7A**). RVLM activation near the vasomotor frequency extracted power from vaso-oscillators, while activation at low frequencies resulted in negative extracted power values, reflecting changes to the background component of the stimulated spectra that were not observed with other stimulation modalities (**Figure 7E**).

Because RVLM and vibrissa stimulation elicited similar arteriole response amplitudes, we have the opportunity to probe, in some sense, competition between interoception and exteroception with the RVLM drive representing interoception and the vibrissa response representing exteroception (**Figure 7F**). The response was quantified across the full cortical mantle in terms of either arteriole area locked to the RVLM frequency or arteriole area locked to the vibrissa frequency (**Figure 7G,H**). When the RVLM stimulation frequency was near the resting-state vasomotor frequency (**Figure 7G**; ordinate), the pial network predominantly locks to the RVLM frequency. However, when RVLM stimulation was presented off resonance and the vibrissa stimulation is closer to resonance (**Figure 7G**; abscissa) pial arterioles are no longer locked to input from RVLM and instead lock to the vibrissa stimulation frequency. This phenomenon is apparent when the same experiments are analyzed in terms of area-locked to the vibrissa frequency (**Figure 7H**).

In total, three different competition experiments, i.e., visual versus vibrissa stimulation (**Figure 6C-E**), RVLM versus vibrissa stimulation (**Figure 7F-H**), and fundamental versus second harmonics from visual (**Figure 6F**) or vibrissa (**Figure 6G**) stimulation, show that spectral proximity to the intrinsic resting-state vasomotor frequency is a dominant factor to arteriole entrainment.

### Sensory and medullary neuronal activation alters resting-state vasomotion patterns

We now turn to the spatial characteristics of the ∼0.1 Hz vaso-oscillations and ask if the driven response merely alters, or fundamentally changes, the underlying spatial patterns of vasodynamic traveling waves^46^ (**Figure 1B,C**). The dominant vasomotor spatial activity patterns were analyzed in terms of the leading space mode as a function of temporal frequency (**Figures S7B-E** and **S8**). Magnitude maps report the spatial distribution of coherence in each mode whereas phase maps indicate differences in timing across the mantle. During resting-state, vaso-oscillations are coherent across the pial network and propagate as traveling waves in symmetric lateral to medial, or lateral to occipital, patterns (**Figure S8A**). The total phase difference is approximately *π* radians across the cortical mantle, consistent with ∼ 2 mm/s vasomotor propagation measured previously^46^. During periodic stimulation near the vasomotor frequency, arteriole power and spatial coherence have peaks at the stimulation frequency and at harmonics (**Figure S8B-D**). How does this translate into changes in spatial phase gradients? First, at the scale of different sensory areas, phase gradients within vS1 cortex increase from rest upon vibrissa stimulation, while phase gradients across V1 cortex are unchanged upon stimulation (**Figure S8E;** p = 6.2×10^-5^ and p = 0.55, one and two-tail Wilcoxon rank sum tests, respectively). Next, at the scale of the entire cortical mantle, vibrissa stimulation increased the phase gradient but not the mean direction of wave motion across the cortical mantle relative to resting-state conditions (**Figure S8F**; 15 animals and 81 experiments, p = 1.3×10^-8^ one-tail Wilcoxon rank sum test). In contrast, visual stimulation did not change the phase gradient across the mantle, while optogenetic activation of RVLM decreased the phase gradient (**Figure S8F**; 87 visual stimulation experiments from 12 animals and 60 RVLM activation experiments from 6 animals, p = 0.59 and p = 0.02, two and one-tail Wilcoxon rank sum tests, respectively), consistent with RVLM activation of spatially uniform neuromodulatory systems^57^. All told, ongoing vasodynamics set the overall pattern of arteriole phase gradients, while the timing and intensity of sensory and subcortical inputs modulate the amplitude and timing of the arteriole dynamics. The most dramatic change is when neural rhythms are on resonance with resting-state vasomotor oscillations.

## DISCUSSION

Vasomotor oscillations modulate pial arteriole diameter, and the perfusion of blood to cortex, by 20% per cycle as measured in mice^46,58^. This modulation occurs through ∼ 20 mm vasomotor waves that are synchronized between brain hemispheres via callosal projections^24^. Thus, cerebral vasomotion, and the parameters that determine its entrainment, are relevant to organ-scale perfusion with applications in both observational, e.g., by BOLD fMRI or fNIRS, and interventional studies of brain physiology. While many of the relevant spatiotemporal properties have been identified in simplified or one-dimensional contexts, collective dynamics of the pial network remained unexplored.

We observed differences in vasodynamics between discrete cortical areas, namely vibrissae and visual cortices, across vasodynamic metrics, including oscillation frequency (**Figure 1F,G**), spectral width (**Figure 1H**), extracted power (**Figure 3E**), stimuli-response amplitude (**Figures 4D** and **5D**), and stimuli-response phase (**Figures 4H** and **5I**). Functional depletion of EC K_IR_2.1 revealed that changes are partially dependent on these channels in both regions (**Figure 2E,F**). Despite regional differences, all areas exhibited resonance near the intrinsic vasomotor frequency (**Table S1**), indicating that resonance is a universal property of the pial network. Collectively, these findings suggest a multifaceted importance of resting-state measurements: First, the vasomotor frequency, which can vary by cortical region, determines the frequency at which the vascular representation of neural input is enhanced in terms of both amplitude and space. In the case of solely vibrissae stimulation, arteriole entrainment reaches far beyond the driven cortical area (**Figure 6A**). Second, resting-state vasoactivity facilitates the stimulated response through the extraction of power from off-resonant neighboring arteriole oscillators near the vasomotor frequency (**Figure 3C-E**). This nonlinear effect, which becomes particularly prominent at large amplitudes (**Figure 3F**), leads to enhanced stimulated responses even in the case of altered neurovascular communication (**Figure 3G**).

### Nonlinear vasodynamics and brain imaging

Our experiments reveal that pial arteriole dynamics become increasingly nonlinear at large amplitudes, and generalization across experiments (**Figure 3F**) suggests that this is a property of the arterioles themselves and not individual stimulation modalities. Notably, extraction of energy from off-resonant neighboring oscillators accounts for ∼ 30% of the line response (**Figure 3F**). This behavior is observed over a 20-fold range of evoked-response power under our conditions, with full-field visual stimulation extracting the lowest power and vibrissa and RVLM stimulation leading to the greatest extracted power (**Figures 3F**). These results provide a potential vascular biophysical basis for differences in functional connectivity derived from vasomotor signal amplitudes^59^.

Our observations on the response of cortex relative to the local, underlying resting-state activity have important implications for the use of BOLD fMRI and fNIRS as reporters of neurological activity. One is that linear response theory, used to define a neurovascular impulse response function (IRF)^4,60^, has restricted applicability in light of the nonlinearity of arteriole vasodynamics (**Figure 3F**). While one can always define an IRF under restricted conditions, the IRF will not take on a constant form across brain areas and stimulus amplitudes^6,61^. At best, an IRF is applicable to weak stimuli in a confined area, as seen for V1 cortex (**Figure 4G-I**). A second point is that the sensitivity of BOLD fMRI and fNIRS as reporters of neurological activity would be increased if the rate of presentation of sensory stimuli were matched to that of the underlying resting-state vasomotion. In this way, the vasodynamic response would increase two-fold in amplitude, which implies a factor of two increase in signal-to-noise ratio or a concomitant decrease in averaging. Indeed, recent data from human arteriole inflow-weighted fMRI supports this hypothesis^62^. A related issue is the designation of the boundary between two brain regions, which can depend on the temporal frequency, waveform, and strength of the stimuli used to probe the two regions (**Figure 6E**). Lastly, the impact of subthalamic inputs, which in principle could be measured from the activity of their nuclei^63,64^, modulates the cortical response (**Figure7F-H**).

### Biophysics

Pioneering measurements identified a large (∼ 50 %) reduction in the CBF response to prolonged whisker stimulation in anesthetized mice after EC K_IR_2.1 knockout^34^. With the knowledge of the relevant channel physiology, we build on this work using awake animals and brief (< 1 s) stimuli that better resemble natural stimuli. We find that loss of EC K_IR_2.1 reduces the pial arteriole response to sensory stimulation by 19 % and delays the response peak by 0.27 s (**Figure 2H**). The integrated response decreased modestly, reflecting its prolonged recovery to baseline after EC K_IR_2.1 knockout (**Figure 2G,H**). The difference in response amplitude reduction is quantitative, not qualitative. Past work measured the fractional change in CBF with laser Doppler flowmetry, introducing sensitivity to underlying vascular beds, as compared to Δ*D*⁄〈*D*〉 measured here by optical sectioning. Further the past measurements used anesthetized mice, while the present work uses awake animals with active NVC feedback^65,66^.

Additional blockade of HCN channels decreases the periodic sensory response by 45-65% compared to K_IR_2.1 knockout alone (**Figure 3G**). The required inhibition of hyper- and re-polarization channels highlights the mechanistic redundancy underlying neurovascular responses^27^. During resting-state, vaso-oscillations were reduced by dual-inhibition but not by EC K_IR_2.1 knockout alone (**Figure 2B,D,E**), suggesting that inhibition of membrane repolarization via HCN blockade may be effective in downregulating vasomotion. Of note, both K_IR_2.1 knockout and simultaneous K_IR_2.1 knockout and HCN blockade resulted in altered stimulus responses that resemble those observed in disease states such as cerebral amyloid angiopathy (CAA) via BOLD fMRI^67,68^. These results may complicate CAA diagnoses via the BOLD response alone and underscore the critical importance of distinguishing between the neurogenic and myogenic contributions to neurovascular dysfunction.

The finding that the resting state spectral width decreases after EC K_IR_2.1 knockout (**Figure 2F**) supports abstraction of the pial arterioles as oscillators that are perturbed by neural input (**Eq 1**). During forcing, arterioles become entrained after a few cycles (**Figures 3A**, and **5G**), suggesting that they operate within the applicable weakly-coupled regime to be described by the Kuramoto model of weakly coupled oscillators^51,69^. Although the Kuramoto model motivated our experiments, it only describes oscillation phase and not amplitude. Yet vasomotor oscillations are broad-band with variable amplitude (**Figure 1D,E,H)**. A hint for the utility of such dynamics comes from comparison with slow oscillations in the gut. There, isolated segments exhibit relatively pure oscillations^70,71^ with a downward shift in the frequency of isolated segments from the level of the stomach to that of the rectum. In the intact system, the frequency gradient is replaced by a staircase of plateaus of decreasing frequencies. This leads, experimentally^71,72^, numerically^36,73,74^, and mathematically^36^ to a traveling wave from stomach to rectum as well as to pinch-offs in the diameter of the gut at the boundary between segments. We similarly observe staircases of decreasing frequency plateaus from vS1 to V1 cortex, consistent with the dominant lateral to occipital resting-state traveling wave patterns (cf **Figures 1F** and **S8A-D**). In the gut, pinch-off leads to a "chain of sausages" that allows food to be churned and mechanically digested. In the bloodstream, pinch-off (vasospasm) is observed in human radiological images from dysfunctional brains, after subarachnoid hemorrhage^75,76^ and in pial arterioles of cynomolgus monkeys made hypertensive by renal ischemia^77^. While vasomotion was not recorded, we note that the stroke prone spontaneously hypertensive rat is known to exhibit endogenous, almost perfectly sinusoid oscillation^18^. We speculate that brain arterioles in healthy animals exhibit broad-band oscillations to avoid severe diminution of blood flow by pinch-off.

### Modeling beyond Kuramoto

Our two key observations about the physical nature of the arteriole dynamics are the extraction of energy from off-resonant oscillators (**Figure 3F**) and the broad width of the spectral response (**Figure 1D-I**). The extraction implies non-linear coupling, such as associated with coupled phase oscillators or spatially extended oscillatory systems that account for amplitude in addition to variations in phase^51,52,78^. The broad width suggests that the observed spectrum from each arteriole segment results from the collective dynamics of a relatively small population of individual oscillators, each with a slightly different natural frequency^79^. While all oscillators are coupled, presumably via connexins to the endothelium^80^, the coupling strength is sufficiently weak so that only a fraction of oscillators are phase-locked with each other. This viewpoint may provide a start to connect the nonlinear arteriole dynamics identified here to an improved inference of neural activity from vaso- and hemo-dynamics.

### Connection to the human condition

In humans, ∼ 0.1 Hz activity dominates resting-state BOLD variability^13^ and vasomotor dynamics, including symmetrically propagating vasomotor waves (**Figure S8**), may explain aspects of BOLD connectivity that are unaccounted for by neuroanatomy^81,82^. Unlike conventional venous-dominated BOLD fMRI, emerging approaches utilizing blood volume and blood velocity weighted signals provide sensitivity to vasomotor artery dynamics in awake humans^62,83^. Post-stimulus ∼ 0.1 Hz ringing^84,85^ revealed by these techniques, and prominent ∼ 0.1 Hz hemodynamic oscillations in surgical patients^86^, suggest that ∼ 10 s vasomotor timescales are relevant in human cerebral arteries.

Vasomotion’s biophysical characteristics pertain to theories of driven brain-waste clearance, including the intramural periarterial drainage pathway (IPAD)^87^ and the glymphatic system^88^. Experiments indicate that driven 0.1 Hz vaso-activity can promote both interstitial fluid movement^89^ and movement of proteins along arteriole walls^90^. Current experimental study of the vascular driving forces enabling the glymphatic system implicate heartbeat^91^ and ultra-low frequency^89^, i.e., ∼ 0.02 Hz, activity. Intrinsic vasomotion is an unexplored intermediary which, compared to heartbeat pulsations, exhibits larger dilation amplitudes, shorter wavelengths, and slower propagation speeds; all differences are by one-to-two orders of magnitude^46^. These features may be especially advantageous in the low Reynolds number environment of the brain’s perivascular spaces. Interventions aimed to drive large-amplitude arteriole dilations to clear brain waste^90^ may be aided by targeting the ∼ 0.1 Hz arteriole resonance (**Table S1**). Further benefit may be gained by altering neurovascular communication to drive spectrally narrowed vasomotion (**Figure 2D,F**).

## LIMITATIONS OF THE STUDY

Simplifying assumptions used in our analyses, including those of stationarity and steady-state dynamics, make it unclear if the large variability observed in the data is completely biological or due to analysis methods. Additionally, measurements of neural activity, via jRGECO in the widefield system, were inherently biased toward excitatory neural processes in superficial layers of cortex. Select findings indicate the involvement of unaccounted for neural mechanisms. First, visual and vibrissa stimulation exhibited comparable peak excitatory neural responses (**Figures 4C** and **5C**), yet vibrissa stimulation elicited a peak arteriole response that was 3.4 times larger than visual stimulation (**Figures 4D** and **5D**). Second, vibrissa and RVLM stimulation produced constriction-biased responses during select low-frequency trials (**Figures 5G** and **S7A**). Given the net-constrictive effect of noradrenaline^60^, and the extensive projection pattern of RVLM to locus coeruleus^57^, it is possible that in some cases noradrenaline release contributes a constrictive nonlinearity. Finally, while K_IR_2.1 knockout and HCN block via ivabradine reported here are not expected to affect the systemic cardiovascular system^34,92^, local loss of channel function could potentially cause a change in autoregulatory state.

## STAR METHODS

### KEY RESOURCES TABLE

| REAGENT or RESOURCE | SOURCE | IDENTIFIER |
|---|---|---|
| Chemicals, peptides, and recombinant proteins |  |  |
| Isoflurane | Henry Schein | Cat no. 1182097 |
| Buprenorphine Hydrochloride, injection | Par Pharmaceutical | Cat no. 110483373347 |
| Cyanoacrylate glue, Loctite 401, Loctite 4014 | MacMaster-Carr | Loctite 401<br>Loctite 4014 |
| Cy5.5 – dextran (2MDa) | Self-synthesized | Liu et al., 2019 |
| Experimental models: Organisms/strains |  |  |
| Mouse: B6.Cg-Tg(Acta2-GCaMP8.1/mVermilion) B34-4Mik/J | Jackson Laboratories | # 032887 |
| Mouse: Tg(Thy1-jRGECO1a)GP8.20Dkim/J | Jackson Laboratories | # 030525 |
| Mouse: Tg(Cdh5-cre/ERT2)1Rha | Taconic Biosciences | Soresen et al.<sup>93</sup> |
| Mouse: C57BL/6J-*Kcnj2*<sup>*em1flox*</sup>/Cya | Mark Nelson | Sonkusare et al.<sup>34,94</sup> |
| B6.FVB(Cg)-Tg(Dbh-cre)KH212Gsat/Mmucd | MMRRC | MMRRC:036778-UCD |
| Software and algorithms |  |  |
| MATLAB | MathWorks |  |
| ScanImage | MBF Bioscience |  |
| Chronux | Chronux.org |  |
| MicroManager 2 | Micromanager.org |  |
| Photoshop 2022 | Adobe Inc. |  |
| Illustrator 2022 | Adobe Inc. |  |
| Other |  |  |
| PowerLab | AD Instruments | Cat no. 16/35 |
| Prime95B | Photometrics | Cat no. Prime95B |
| Two photon microscope with adaptive optics | Thorlabs and others | Liu et al.<sup>95</sup> |
| SteREO Discovery.V8 | Zeiss | Broggini, Duckworth et al.<sup>46</sup> |

### RESOURCE AVAILABILITY

#### Lead contact

Further information and requests for resources and reagents should be directed to and will be fulfilled by Prof. David Kleinfeld.

##### Materials availability

N/A

#### Data and code availability

The datasets supporting the current study, and an associated "read me" file, will be available at https://www.dandiarchive.org/ upon publication. All code will be available upon publication at https://github.com/JDuckworth1.

### METHOD DETAILS

#### Experimental subjects

All animal procedures were approved by the Institutional Animal Care and Use Committee at the University of California, San Diego (UCSD) and were performed under the guidance of the UC San Diego Center for Animal Resources and Education.

#### Animals

The B6.Cg-Tg(Acta2-GCaMP8.1/mVermilion)B34-4Mik/J (Jax 032887) smooth muscle expressing GCaMP8.1 animals were crossed with STOCK Tg(Thy1-jRGECO1a)GP8.20Dkim/J (Jax# 030525) neuronal jRGECO1a expressing animals. For RVLM experiments, the smooth muscle expressing GCaMP8.1 mice were crossed with dβh-cre (MMMRC B6.FVB(Cg)-Tg(Dbh-cre)KH212Gsat/Mmucd) expressing animals. Prior to surgery the animals were housed one to five per cage with a 07:00 to 19:00 light-dark cycle. Variability across behavioral and physiological traits are not significantly greater in female rodents, including across the estrous cycle, than male rodents.^96,97^

#### Surgery

Adult mice of both sexes were anesthetized with 4 % (v/v) isoflurane in oxygen for induction and 1 – 3 % (v/v) for maintenance. Reflexes and breathing were visually monitored through the entire surgical procedure to ensure a deep plane of anesthesia. Body temperature was maintained at 37 C with a heating pad with feedback regulation (FHC 40-90-8D). The animal was placed in a stereotaxic frame, the scalp was sterilely removed, the periosteum on the parietal and occipital plates was gently and thoroughly removed, skull sutures were covered with low viscosity cyanoacrylate glue (Loctite 4014) to reinforce stability between skull plates, and an approximately 10 mm by 10 mm region of skull over cortex was thinned with a 250 µm diameter drill bur coupled to a low vibration drill (Osada EXL-M40) to form a transcranial window. The thinned bone was dried and covered with cyanoacrylate glue (Loctite 401) and a number 0 glass coverslip. A titanium implant was glued onto the skull posterior to lambda for head-fixation. The remaining exposed bone and the implant were covered with cyanoacrylate glue and dental cement (Lang, Ortho-Jet^TM^) to increase stability. Buprenorphine hydrochloride (Buprenex, Reckitt Benckiser Pharmaceuticals) was provided subcutaneously for analgesia (2 mg per kg body weight) as the animals recovered from surgery.

#### Post-surgical preparation

After a minimum of 48 hours post-surgery, mice were handled and trained to head fixation in sessions of 20 minutes to 3 hours per day for 2 days, and then trained an additional 1 to 3 days at 3 hours per day for awake head fixation under the microscope.

#### Widefield imaging

A Zeiss SteREO Discovery.V8 was equipped with a custom filter cube (KSC 295-CUSTOM, Kramer Scientific LLC) carrying a reflective mirror and ZET488/561m emission filter (Chroma). Dual color imaging is achieved using interlaced activation of two LED at ∼10 Hz (ThorLabs M470L3 with 470 nm center wavelength and Chroma ET470/40x filter; ThorLabs SOLIS-4C with 565 nm center wavelength and Semrock FF01-560/14-25 filter) coupled to a liquid lightguide. Image acquisition was performed with a Prime95B 1200×1200 (Teledyne Photometrics) and MicroManager software. The microscope configuration can be found online: https://www.fpbase.org/microscope/spJ5csFwGSy62EDQkoJPRd/?f=filter-ex-408,filter-bs-2989,filter-em-2&p=gcamp6f_in-presence-of-ca2&l=thorlabs-m470l4-led&d=photometrics-prime-95b.

#### Sensory stimulation and experiment design

Stimulation was timed with custom written MATLAB code that controlled a USB-6343 board (National Instruments). All widefield experiments were performed with the animals awake and head fixed. The first 250-500 s were recorded without stimulation and were used for vasomotor frequency quantification. The second 500 s was recorded during periodic stimulation, with frequency ranging from 0.02 Hz to 0.4 Hz. Tactile vibrissa stimulation was delivered unilaterally or bilaterally and consisted of three 10 ms air puffs, separated by 150 ms. Visual stimulation was delivered unilaterally by a fiber coupled 448 nm LED (ThorLabs M445L3), which delivered 50 Hz pulses at 50 percent duty cycle for 2 s. Visual grating stimulation was performed using an LCD screen (Dell P2210) with the dimensions 18.7 inches by 11.7 inches and 1680 x 1050 resolution. The horizontal grating was presented with 1.5 cycles / sec to one eye of the animal using custom written MATLAB code utilizing the Phychtoolbox package. Two-photon imaging experiments were performed with awake and head fixed animals and consisted of 300 s without stimulation followed by 300 s of 0.1 Hz air puffs directed unilaterally to the vibrissae. Here we used 10 ms air puffs presented at 20 Hz for 0.5 s.

#### Optogenetic RVLM activation

Dβh-cre animals crossed with SMC GCaMP8.1 were used for these experiments. Three weeks prior to thin-skull surgery, animals were injected (70 nl at 5 nl/min using a Nanoject injector) with a cre-dependent ChRmine virus (AAV-8-Ef1a-DIO-ChRmine-eYFP-WPRE, obtained from Stanford viral core) resulting in opsin expression localized to catecholaminergic dopamine-β-hydroxylase positive neurons in the RVLM. The expression was confirmed postmortem with co-labelling with anti-Tyrosine Hydroxylase antibody (Sigma-Aldrich, AB152) and counterstained with Neurotrace blue (ThermoFisher, N21479) (**Figures 7A** and **S6**). Histological sections of thickness 30 µm were mounted on glass slides, coverslipped with fluoromount (Invitrogen, Catalog number 00-4958-02) and scanned on Zeiss axioscan. Optogenetic stimulation was delivered through the ear with a custom built 630 nm LED illuminator, using 50 Hz pulses at 50 percent duty cycle for 2 s. For each animal, a set of experiments with increasing LED power were performed to identify animals that exhibited a SMC calcium response to optogenetic stimuli (**Figure S6**). Reported LED power is the DC power measured using a Coherent FieldMate power meter. Animals that exhibited response line-power greater than 0.1, when normalized by the experiment’s total power, were classified as responders and included in the results, with LED power of 100 mW (**Figures 7C-E**).

#### Induction of EC K_IR_2.1 knockout and ivabradine administration

A first set of experiments were performed using double-crossed animals that express tamoxifen-inducible Cre under the vascular endothelial cadherin promoter (Cdh5-CreERT2)^93^, and a “floxed” version of the kcnj2 gene (i.e., kcnj2 flanked by loxP sequences)^34,94^. Control mice expressed cdh5-CreERT2 but not floxed-kcnj2 and underwent identical tamoxifen treatment as the experimental group.

Following pre-tamoxifen treatment imaging sessions, recombination was induced by retro-orbitally injecting 10 mg/kg tamoxifen (Sigma-Aldrich #T5648) solubilized in ethoxylated castor oil for 5 consecutive days^98^. A minimum period of 7 days surpassed, after the final day of tamoxifen treatment, before follow-up imaging to allow for recombination and protein turnover. Ivabradine (Sigma-Aldrich #SML0281) was dissolved in 1x PBS and injected retro-orbitally. We used 1 mg/kg for widefield Ca^2+^ imaging experiments and 10 mg/kg for two-photon measurements. We waited 20 minutes following injection and cessation of isoflurane before imaging to allow for recovery of arteriole reactivity. Data was collected for up to 2 hours following ivabradine injection; longer times resulted in diminished effects. Using two-photon microscopy with resonant scanning, we imaged the same pial arterioles in awake mice before and after K_IR_2.1^95^ knockout and ivabradine administration. As these were pair-wise measurements, we used one-tail Wilcoxon signed rank tests unless otherwise noted.

#### Data Analysis

##### Mask creation

As described previously^46^, to create each trial’s pial arteriole mask, we took the standard deviation over time of the trial’s first 1,250 diffusion filtered GCaMP8.1 fluorescence images. The result was normalized and contrast-enhanced using MATLAB’s "adapthisteq" function. Adobe Photoshop was used to sharpen, threshold, and manually connect pial arteriole segments that fell below the threshold. The neural mask was manually defined inside the thin-skull window edge, excluding the pial arteriole mask. The pial arteriole mask was used to define a vessel graph using procedures previously described^99^.

#### dF/F Calculation and definition of the vasomotor frequency

Changes in fluorescence, dF/F_x,t_, were calculated from masked images as described previously^46^. To calculate resting state spectra and perform line component analyses, we first performed a space-time singular value decomposition (SVD) on vascular and neuronal data, keeping the first T/2 and T/10 modes respectively, where T is the trial length in samples. We defined each trial’s vasomotor frequency using the baseline time period’s SMC-GCaMP dF/F_x,t_ multitaper spectrum, with half-bandwidth = 0.02 Hz and 19 tapers. Spectra from all pixels were averaged across space to obtain the baseline vascular spectrum. The frequency that maximized the baseline spectrum’s power within a user-defined 0.05 – 0.25 Hz range was defined as the vasomotor frequency and used for subsequent analyses. For results restricted to primary sensory cortices (**Figures 3E** and **6E**) we used baseline spectra from pixels within the relevant sensory cortex, as opposed to all pixels, to define the vasomotor peak.

#### Spectral analysis

All spectral quantities were calculated in MATLAB using the multitaper method with Slepian tapers. Calculations followed normalization conventions provided by Persival and Walden^50^ and used some helper functions from the Chronux toolbox. The number of tapers was 2TW-1 where T is the acquisition length and W is the Slepian taper half-bandwidth.

#### Quantification of resting-state vaso-oscillations

Resting-state arteriole oscillations were characterized in terms of their spectral properties, namely their peak spectral power, frequency at peak power, spectral width *BW*, and difference in frequency δ*f*. Multitaper spectra were first calculated from each arteriole segment’s time series and temporal derivative time series; taking the temporal derivative served to remove ∼ 1/f^2^ spectral background. For each experiment, these spectra were plotted, and manual frequency bounds were selected to isolate ∼ 0.1 Hz peaks. Peak spectral power and frequency at peak power were taken from the raw multitaper spectra, at the point that maximized spectral power within the user-defined frequency bounds. Frequency differences, δ*f*, were the maximum consecutive change in peak frequency, taken along each arteriole segment. Here only arteriole segments longer than 0.5 mm were considered, and frequency profiles were 5-point median filtered before calculating the maximum consecutive change. Spectral widths were the full width at half maximum of the derivative spectra within the user-defined frequency bounds. Here baseline was defined to be the lowest point on either side of the bounded peak. Arteriole segments whose spectra did not cross the half-maximum value on both sides of the peak were removed from the analysis.

#### Quantification of stimulated responses

Thomson’s test for periodicity in colored noise was used to quantify the vascular and neuronal response to stimulation^50^. We modeled activity as background noise plus sinusoids, **Eq 1**, or equivalently:

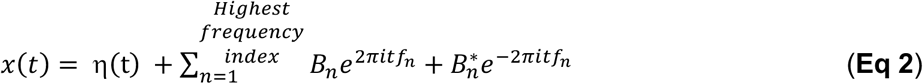

where 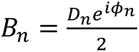 is the complex line amplitude for frequency of interest *n.* Multitaper Fourier transforms of *x*(*t*) were used to calculate 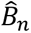, the best estimate line amplitudes using a regression across tapers^50^. The 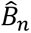 were then used to calculate F-values to assess each line component’s significance relative to uncorrelated noise. Critical F-values were calculated using the inverse F-cumulative distribution, with 2 and 2K-2 degrees of freedom and p = 1/N where K is the number of tapers and N was the number of time points in the experiment. This procedure was performed at every pixel (or arteriole segment) in the field of view.

Residual spectra were calculated by removing significant line components from the total activity in the time domain, as implemented in Chronux, and calculating the multitaper spectrum of the result, denoted *S*_η_(*f*). Spectra were then plotted as

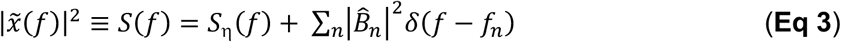

for *n* significant line components as determined by the F-test. The second term was plotted as a delta functional with height 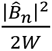 where *W* is the half-bandwidth of the Slepian tapers, chose to be 0.02 Hz unless noted otherwise. Responses were reported using the line-power at the frequency of interest *n*, 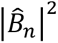, with units 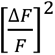. In various cases, 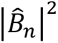 was also plotted pixel-wise onto the neuron and arteriole masks or averaged across the stimulated cortical region (V1 or vS1 cortices) for summary plots. The reported phase responses are the angle of 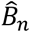, averaged across the stimulated cortical region using a circular mean. For SMC Ca^2+^ data, we plot the phase of the sign-inverted GCaMP8.1 signal. Given that the false positive rate for detecting a line component is 1/N, experiments that did not exhibit more than N_pixels_/N significant line components in the stimulated region (where N_pixels_ is the number of pixels in the cortical region) were deemed to have no significant response and were plotted as open circles on summary plots. These experiments were excluded from phase plots. This criterion was also applied to analyses across the full cortical mantle.

Line-power was used to quantify the spatial extent of entrainment. We defined a pixel’s locked frequency as the frequency at maximum line-power 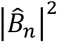. If this frequency was within *f_stimulation_* ± 0.005 Hz, we considered it entrained to the stimulation. Areas were calculated as the fraction of pixels within the stimulated hemisphere/hemispheres (**Figures 6A,B**), stimulated cortical region (V1; **Figure 6C-E**), or full cortical mantle (**Figures 7G,H**) that were entrained to the relevant stimuli. For resting state periods, we did not perform a test for periodicity and instead plotted each pixel’s frequency at maximum spectral power (**Figures 1F** and **7B, left**).

Extracted power was calculated by subtracting the residual spectrum, *S*_η_(*f*) from the experiment’s resting-state spectrum. The resulting difference spectrum was integrated at and around the stimulation frequency, with bounds that were set by the closest zero-crossing points in the difference spectrum. Bounds were limited to be at most ± 0.02 Hz from the stimulation frequency.

Sensory cortices (V1 and vS1 cortices) were located using the Allen Atlas common coordinate frame^100^. For each experiment, the top-down atlas projection was scaled according to the image resolution and manually registered using the leading modes of a space-time SVD on the trial’s neural data during stimulation. These SVD modes reliably exhibited localized barrel and visual cortex activity, regardless of the experiment type. For experiments without neural data, we used manually drawn sensory regions based on Paxinos stereotaxic coordinates^101^.

#### Space frequency analysis

A space-frequency SVD was used to visualize the dominant vascular dynamics present in the data^102^. We first calculated the Fourier transform of each pixel’s tapered time series, forming the complex matrix V(s,f,k). For each frequency, f = f_0_, we calculated the SVD of V(s,f_0_,k). The relative size of the first singular value provides an estimate of spatial coherence. From here, frequencies of interest were chosen from peaks in the global coherence spectrum. Maps (**Figure S8A-D**) are the magnitude and phase of the first spatial mode at each specified frequency. The magnitude map indicates the spatial distribution of coherence, and the phase map indicates the direction of propagation within the vicinity of each specified frequency. Here we defined phase such that the relative delay between two locations increases with increasing phase.

#### Curve fitting

Quantile regression was used to estimate 50^th^ percentile estimates using second order Epanechnikov kernels and the R “np” library for nonparametric kernel smoothing^103^. Fixed x-bandwidths of 0.01 Hz and 0.02 Hz were used for fundamental 2^nd^ harmonic line-power data, respectively. Adaptive nearest-neighbor bandwidth determination was used for extracted power and entrained-area plots. Percentiles were fit with splines using MATLAB’s “spap2” function before plotting. 95 % confidence intervals for the 50^th^ percentile curves were found using a bootstrap procedure in which 10,000 replicate 50^th^ percentile curves were calculated after sampling all data points with replacement.

To compare on to off-peak enhancement, we compared the median response value at peak to their value at 0.04 Hz; this frequency was chosen conservatively as it is off-resonance but not beyond the ∼ 0.2 Hz high frequency limit of the arteriole response. For visual comparison (**Figures 4D,F** and **5D,F**) we scaled neural median curves by (A_0.04_/N_0.04_) before overlaying them on arteriole response curves. Here A_0.04_ is the arteriole median value at 0.04 Hz and N_0.04_ is the neural median value at 0.04 Hz. For 2^nd^ harmonic responses, 0.18 Hz was used as a reference instead of 0.04 Hz.

Extracted power versus line-power data were fit to a linear model on the log-log scale, i.e. *y* = *ax^b^* in linear coordinates. Before fitting, points were excluded and plotted as open circles if their Cook’s distance (found using MATLAB’s “fitlm”) exceeded three times the dataset’s mean Cook’s distance. For the summary across datasets (**Figure 3F**), the slope and intercept were first left as free parameters, yielding a slope close to one (*b* = 0.93 ± 0.07, mean ± SE). A second model was then fit with slope fixed at one, yielding *a* = 0.3 ± 0.02, mean ± bootstrap SE. For individual datasets (**Figure S3A,B**), we also report the R^2^ of linear fits to subsets of the data, to estimate a possible transition point. Here each R^2^ corresponds to a fit that includes all points greater than or equal to the reported breakpoint. Black lines report the linear fit for which R^2^ was maximized.

For visual and vibrissa stimulation, we fit the line-power to the following model of a driven harmonic oscillator:

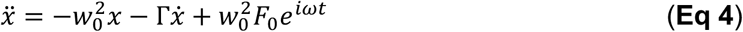

where F_o_ scales the input amplitude and is the value of x at zero frequency. The Lorentzian-type amplitude solution for the steady-state response *x*(*t*) = *Ae^i^*^(^*^ωt^*^+^*^φ^*^)^ is:

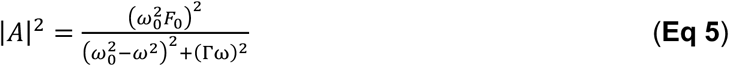

with phase, which we fit for the case of visual stimulation:

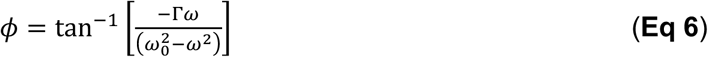

Here, fits and 95 % bootstrap confidence intervals were calculated in MATLAB using nonlinear regression.

For two-dimensional competition experiment summaries (**Figures 6E** and **7G,H**), data were logit transformed and fit to a quadratic surface using MATLAB’s “fit” function with “poly22” fit type.

## Acknowledgments

We extend our gratitude to Randy L. Buckner, Anna Devor, Scott Earley, Prakash Kara and Jonathan Polimeni for discussions, and to Beth Friedman for detailed comments on drafts of the manuscript. This work was supported by the NIH BRAIN initiative grant U19 NS123717 as well as NIH grants R35 HL140027, R01 NS110656, R35 NS09726, RF1 NS128963. TB was additionally supported by the SNF Grants P2SKP3_164948 and P300PA_177804, DK was additionally supported by the Dr. George Feher Experimental Biophysics Endowed Chair, and MTN was additionally supported by the Leducq Foundation Transatlantic Network of Excellence (International Network of Excellence on Brain Endothelium: A Nexus for Cerebral Small Vessel Disease 22CVD01 BRENDA), and the Totman Medical Research Trust.

## Author Contributions

D.K. conceived the study. T.B., J. D., D.K. and M. V. designed the experiments. T.B., K.C. and J.D., performed all experiments and analyzed data with input from S.I., D.K., M.S.-P. and M.V. J.D. and D.K. wrote the manuscript with input from all authors. D.K. attended to the university rules and forms that govern environmental health and safety, including the ethical use of animals as well as the use of chemicals, controlled substances, hazardous substances, lasers, and viral agents.

