## Supplemental Table and Figures for "Neurons entrain and shape an intrinsic arteriole rhythm that distributes blood across the cortical mantle"

#### Table S1. Neural and arteriole resonance parameters

| Drive | | Neurons | | Arterioles | | | Quality (Q)# | | Peak-to-reference ratio^ | | |
|---|---|---|---|---|---|---|---|---|---|---|---|
| | | $f_{vaso}$ (Hz) | FWHM (Hz) | $f_{vaso}$ (Hz) | FWHM | $R^2$ | Neurons | Arterioles | Neurons | Arterioles | Normalized arteriole response |
| **Visual** | **f** | 0.13 | 0.09 | 0.11 | 0.05 | 0.35 | 1.4 | 2.0 | 2.8 | 3.9 | 1.9 |
| | **2f** | 0.13 | 0.17 | 0.10 | 0.09 | 0.73 | 0.7 | 1.0 | 1.1 | 3.7 | 3.2 |
| **Vibrissa** | **f** | 0.16 | 0.13 | 0.13 | 0.03 | 0.34 | 1.3 | 3.6 | 2.2 | 5.3 | 2.3 |
| | **2f** | 0.14 | 0.10 | 0.12 | 0.09 | 0.52 | 1.5 | 1.3 | 1.2 | 3.5 | 4.9 |
| **RVLM** | **f** | - | - | 0.11 | 0.06 | - | - | 1.9 | - | 6.6 | - |
| **Average enhancement** | | | | | | | **1.2** | **2.0** | **1.8** | **4.6** | **3.1** |

### Q ≡ $f_{vaso}$/FWHM. ^ Calculated relative to line power at 0.04 Hz (fundamental) or 0.18 Hz (2nd harmonic)

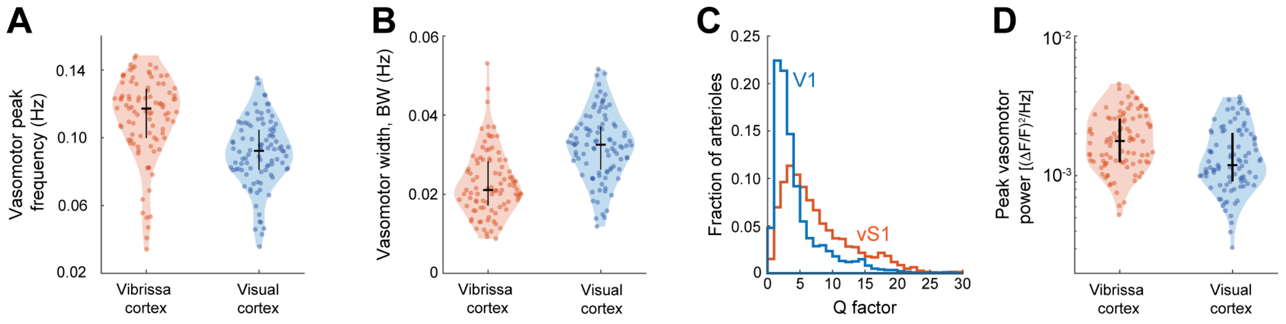


**Figure S1. Vasomotor oscillations differ between vibrissa and visual cortex, related to Figure 1**

(**A**) Peak vasomotor frequency averaged over pial arterioles in vibrissa (vS1; orange) and visual (V1; blue) cortex of awake mice. One estimate per area was calculated for each experiment. Bars denote the 25th, 50th, and 75th percentiles. (**B**) Spectral width, BW, of the vasomotor peak for the same areas as in (**A**).
(**C**) Distribution of the resting-state spectral peak Q factor for arterioles in vibrissa and visual primary sensory cortices. Distributions include data pooled across arteriole segments and experiments (95 experiments, 103,199 and 88,713 total arteriole segments from V1 and vS1 respectively). (**D**) Peak vasomotor power, for vibrissa and visual primary sensory cortices, plotted as in (**A**).

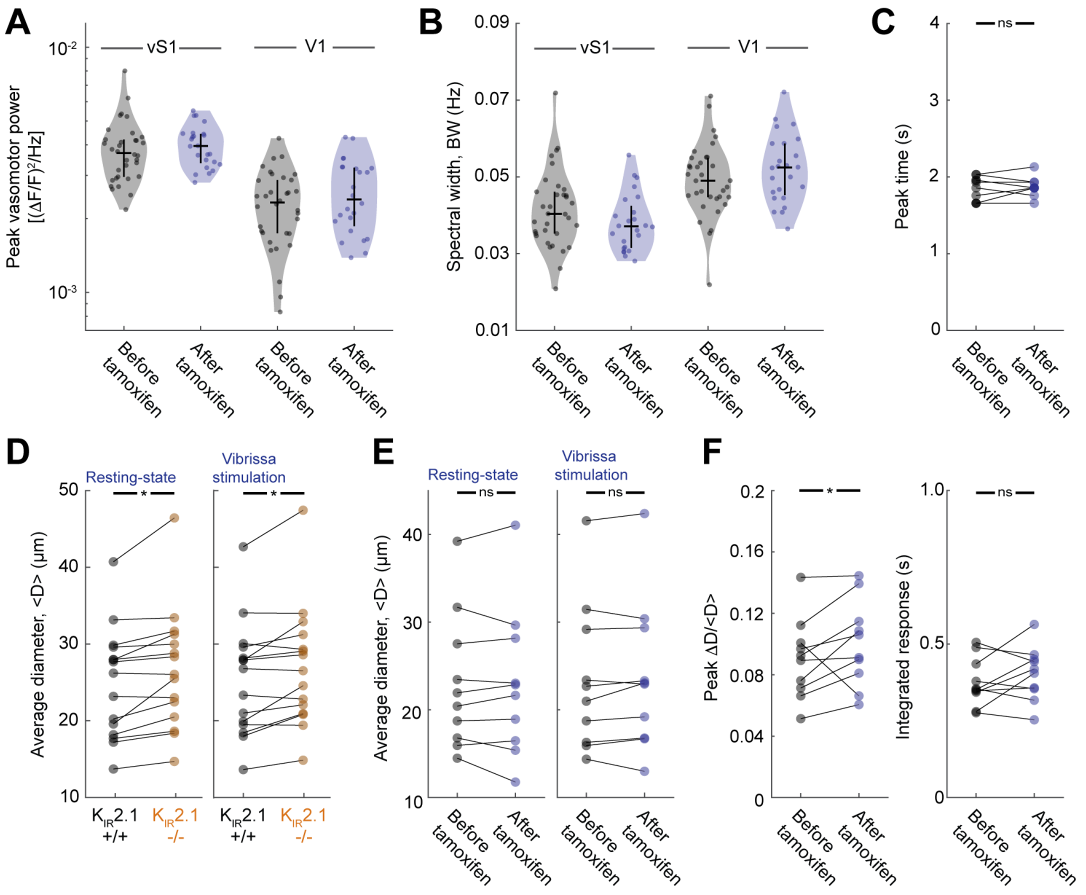


**Figure S2. Vaso-oscillations are unaffected by tamoxifen administration in control animals, related to Figure 2**

(**A**) Peak vasomotor power in vS1 and V1 before and after tamoxifen administration in control animals. Points represent the vasomotor power, averaged across arterioles in the given sensory area; one estimate per area was calculated for each experiment. Bars denote the 25th, 50th, and 75th percentiles.
(**B**) Spectral with, BW (FWHM), of the vasomotor peak for the same datasets shown in (A).
(**C**) Time to peak diameter change, calculated from the stimulus-triggered average trace, during 0.1 Hz vibrissa stimulation in control animals. Lines connect measurements from the same arteriole before and after tamoxifen treatment (**D**) Average arteriole diameter, <D>, in the experimental group ($K_{IR}2.1^{fl/fl}$) before and after taxmoxifen treatment during 300s resting-state and 0.1 Hz vibrissa stimulation experiments. Asterisk denotes $p < 0.05$.
(**E**) Average diameter (<D>) before and after tamoxifen treatment in control mice. (**F**) Peak diameter change, ΔD/<D>, and integrated dilation response during 0.1 Hz vibrissa stimulation before and after tamoxifen in control animals.

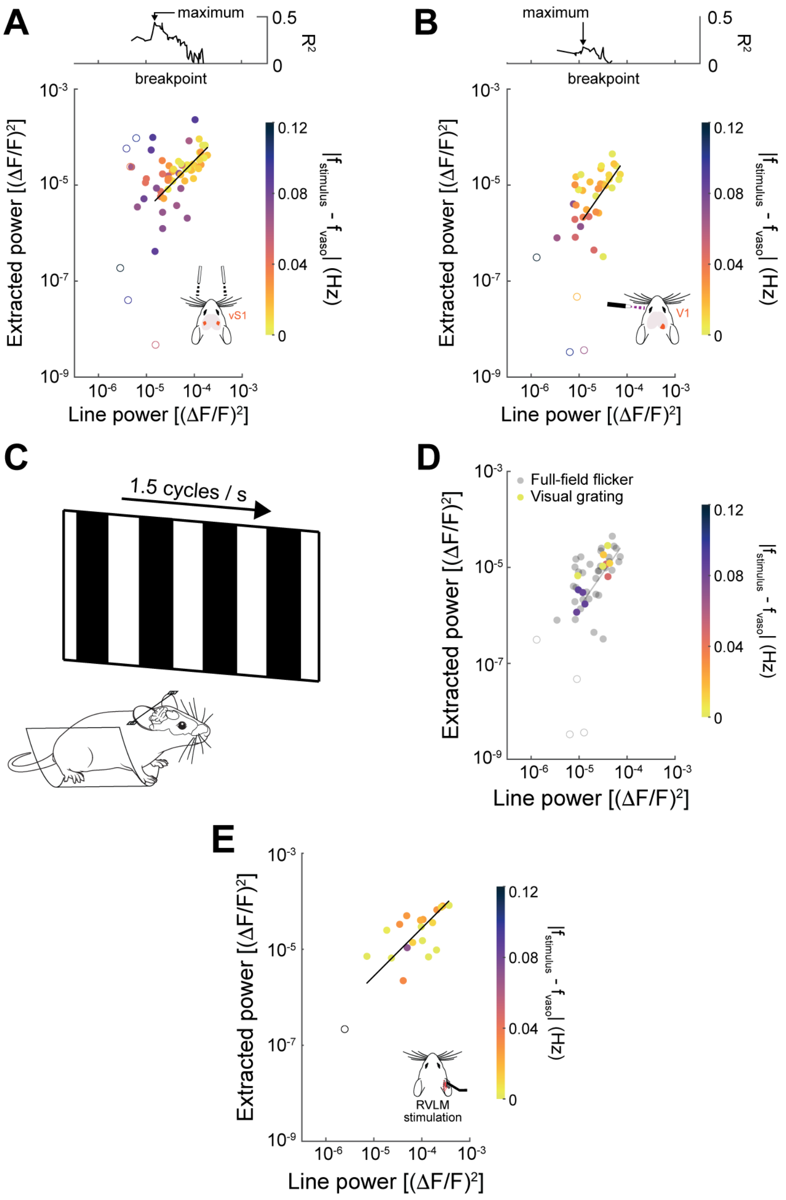


**Figure S3. Resting-state power is systematically extracted during vibrissa, visual, and RVLM stimulation, related to Figure 3**

(**A**) Extracted spectral power versus line-power at the stimulus frequency for arterioles in vS1 during vibrissa stimulation. Open circles were excluded from the linear fit, which had a maximum $R^2 = 0.44$ when only experiments with line power greater than 1.5 x 10-5 ΔF/F were included (log-log slope = 1.0 ± 0.2, intercept = -0.43 ± 0.71; regression estimates ± SE, see **STAR METHODS** for details). Marker color encodes the offset between stimulus and endogenous vasomotor frequency, $|f_{stimulus} - f_{vaso}|$. Top: coefficient of determination, $R^2$, of the piecewise fit as a function of breakpoint; arrow marks the maximum, which defines the threshold used for the fit line. (**B**) As in (**A**) for V1 during full-field visual stimulation, for arterioles overlying V1. A linear fit to only experiments with line power greater than $1.2 \times 10^{-5}$ ΔF/F maximized $R^2$ ($R^2 = 0.32$; log-log slope = 1.4 ± 0.4, intercept = 1.3 ± 2.0, regression estimates ± SE). (**C**) Schematic of the drifting square wave grating stimulus, reversing at 1.5 cycles/s. (**D**) Extracted versus line-power for full-field flicker (gray) and drifting gratings (colored), ploteted as in (**B**).
(**E**) As in (**A**) for arterioles across the cortical mantle during optogenetic stimulation of the RVLM. A linear fit with slope fixed to one yields an intercept = -0.56 ± 0.09 (black line).

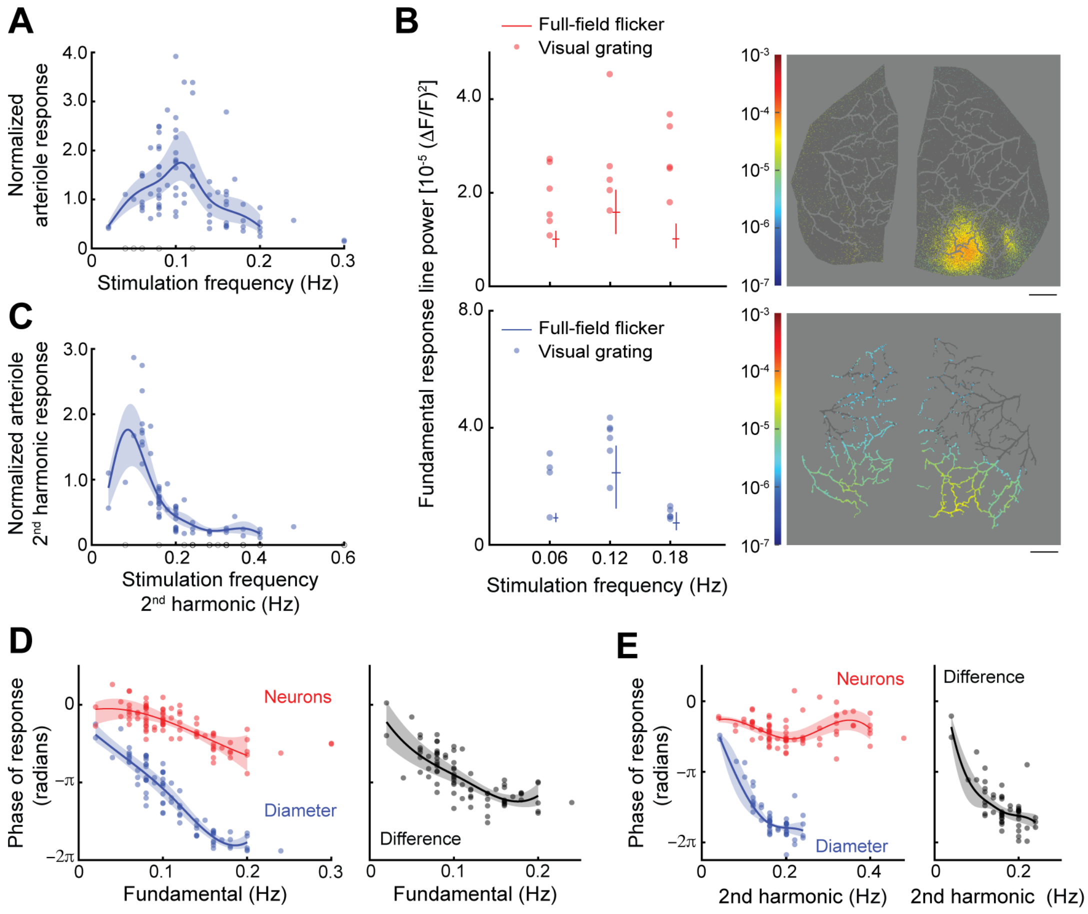


**Figure S4. Normalized arteriole responses to full-field visual stimulation, and visual grating response characterization, related to Figure 4**

(**A**) Arteriole line-power, after normalization by neuronal line-power, as a function of stimulation frequency; line is a kernel-smoothed median and 95% CI band. Open symbols are non responders.
(**B**) Line power at the stimulation frequency during periodic moving visual grating experiments at three frequencies. Solid lines denote $25^{th}$, $50^{th}$, and $75^{th}$ percentiles of the full-field visual stimulation response (**Figure 4B**), for reference. Line-power maps for neural (top right) and arteriole $Ca^{2+}$ (bottom right) during an example 0.18 Hz experiment. (**C**) Arteriole line-power at the stimulation frequency $2^{nd}$ harmonic frequency after experiment-wise normalization by neural $2^{nd}$ harmonic line power during visual stimulation, as plotted in (**A**). (**D**) Phase response during full-field visual stimulation experiments. Diameter phase was calculated using the previously quantified SMC $Ca^{2+}$ - diameter phase relationship[1]. (**E**) Phase response at the stimulation $2^{nd}$ harmonic frequency during full-field visual stimulation experiments, as calculated and plotted in (D).

1. Broggini, T., Duckworth, J., Ji, X., Liu, R., Xia, X., Mächler, P., Shaked, I., Munting, L.P., Iyengar, S., Kotlikoff, M., et al. (2024). Long-wavelength traveling waves of vasomotion modulate the perfusion of cortex. Neuron 112, 2349-2367.

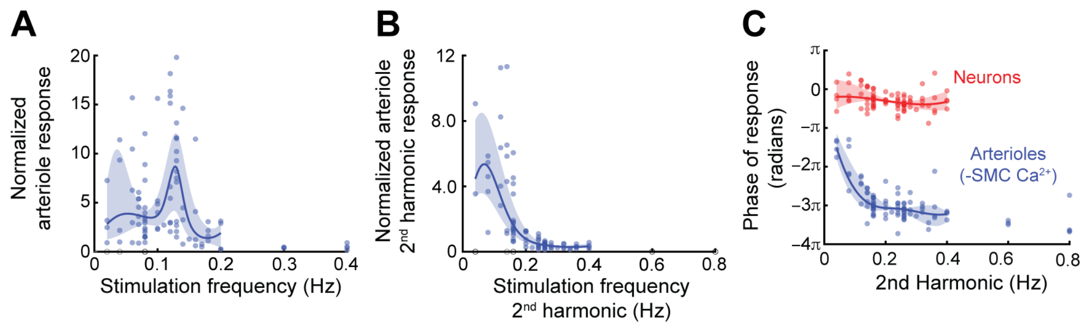


**Figure S5. Normalized arteriole responses and 2nd harmonic response phase during vibrissa stimulation, related to Figure 5**

(**A**) Arteriole line-power after experiment-wise normalization by neural line-power for periodic vibrissa stimulation experiments. Curves denote the median and 95% confidence interval of the median. Open circles denote experiments that did not exhibit significant line power in vS1, in either the arteriole or the neural response. (**B**) Arteriole line-power at the stimulus 2nd harmonic frequency after experiment-wise normalization by neural 2nd harmonic line-power during vibrissa stimulation, plotted as in (**A**). (**C**) Phase of the response at the stimulation 2nd harmonic frequency during vibrissa stimulation for neurons (red) and arterioles reported by SMC $Ca^{2+}$, sign-inverted (blue), versus second-harmonic frequency.

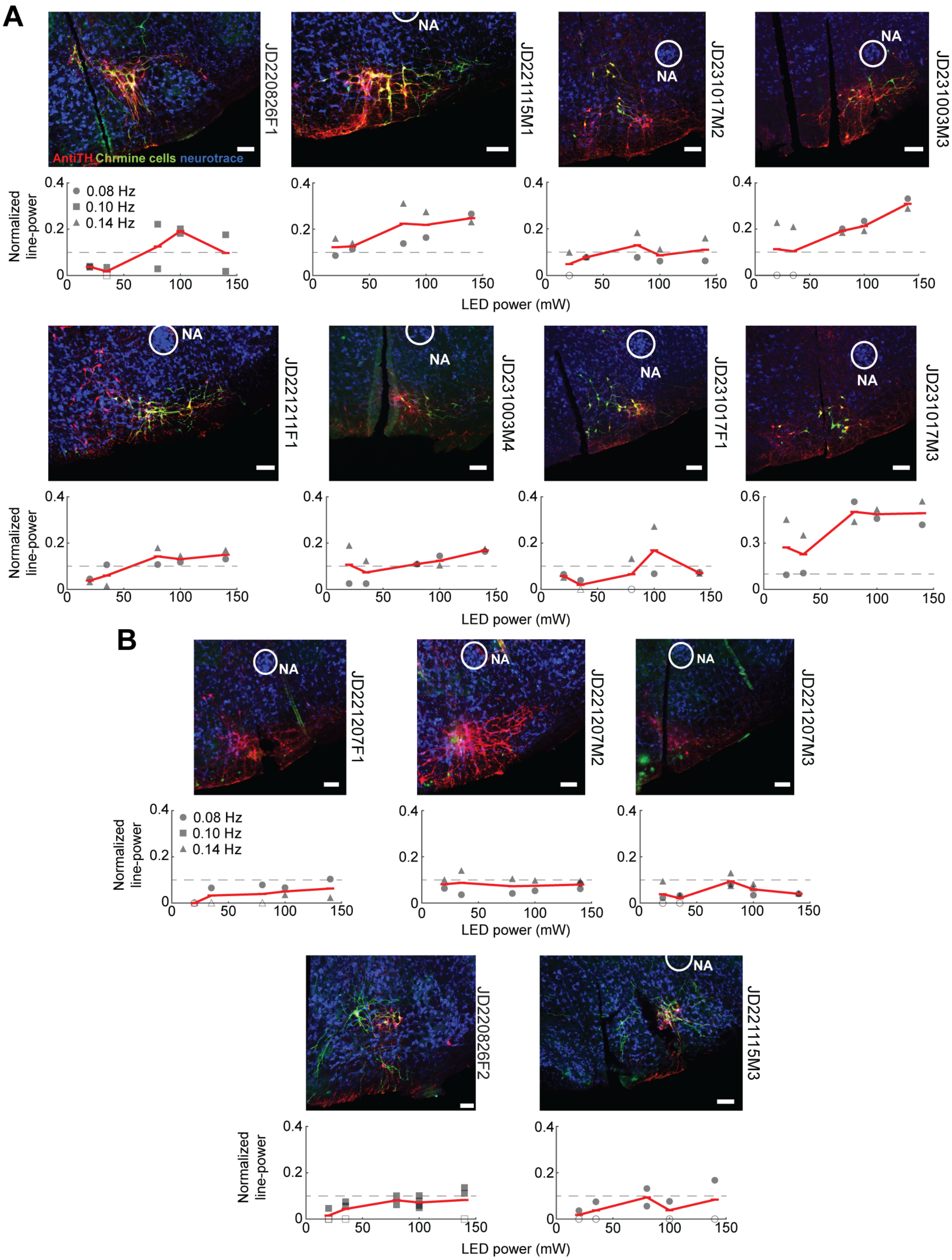


**Figure S6. Histological verification of ChRmine expression in the RVLM combined with arterial response power relative to LED power, related to Figure 7**

(**A**) Mice classified as responders to optogenetic RVLM activation. For each animal: coronal section immunostained for tyrosine hydroxylase (red), ChRmine-expressing cells (green), and NeuroTrace counterstain (blue), with the nucleus ambiguus circled for reference; animal identifier at right. Scale bars, 100 μm. Below each section, normalized line-power of the arteriole response as a function of LED power input at three stimulation frequencies (0.08, 0.10 and 0.14 Hz). Red line is the mean across experiments, dashed line marks the chosen threshold, and open shapes indicate a trial with an insignificant number of line components. Animals exhibiting a mean normalized response at any LED power that surpassed 0.1 were classified as responders. (**B**) As in (**A**) for mice classified as non-responders.

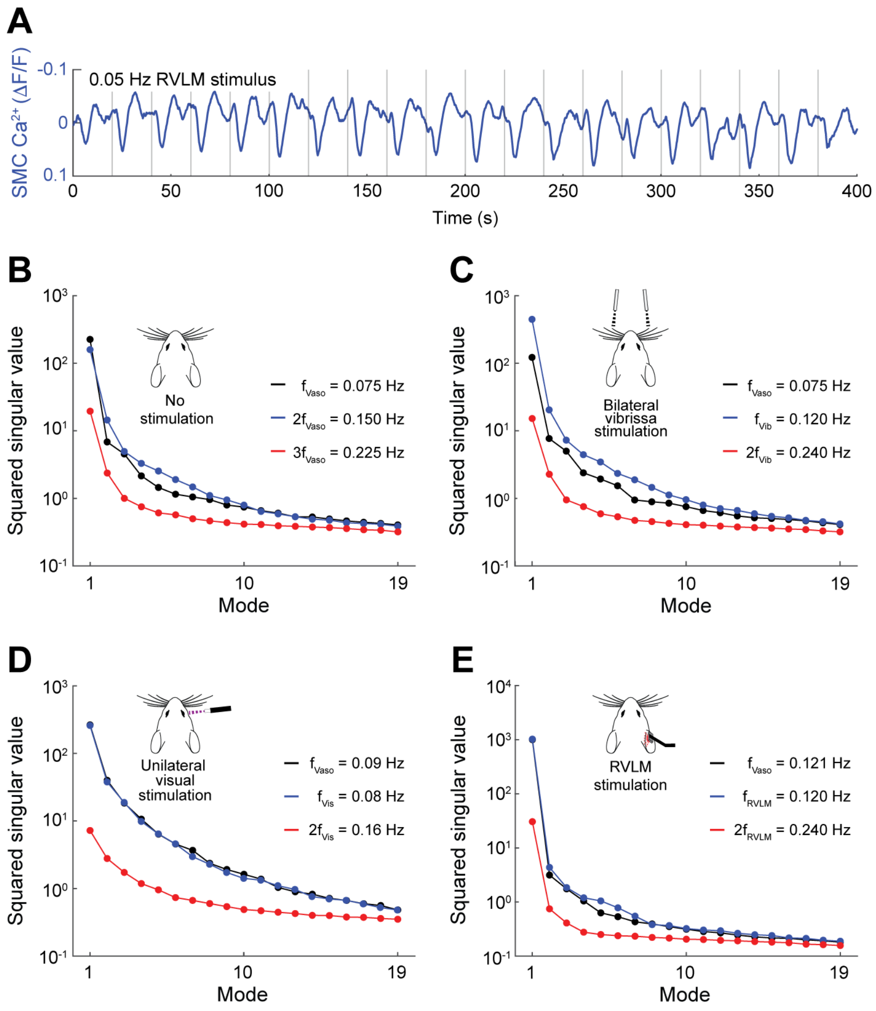


**Figure S7. Constrictive peaks during low-frequency RVLM stimulation, and space-frequency SVD singular value spectra, related to Figure 7**

(**A**) Arteriole SMC $Ca^{2+}$ time series for one example RVLM activation experiment at $f_{RVLM}$ = 0.05 Hz. Gray lines indicate stimuli time points. (**B-E**) Space-frequency SVD singular value spectra corresponding to the analyses presented in **Figure S8A-D**.

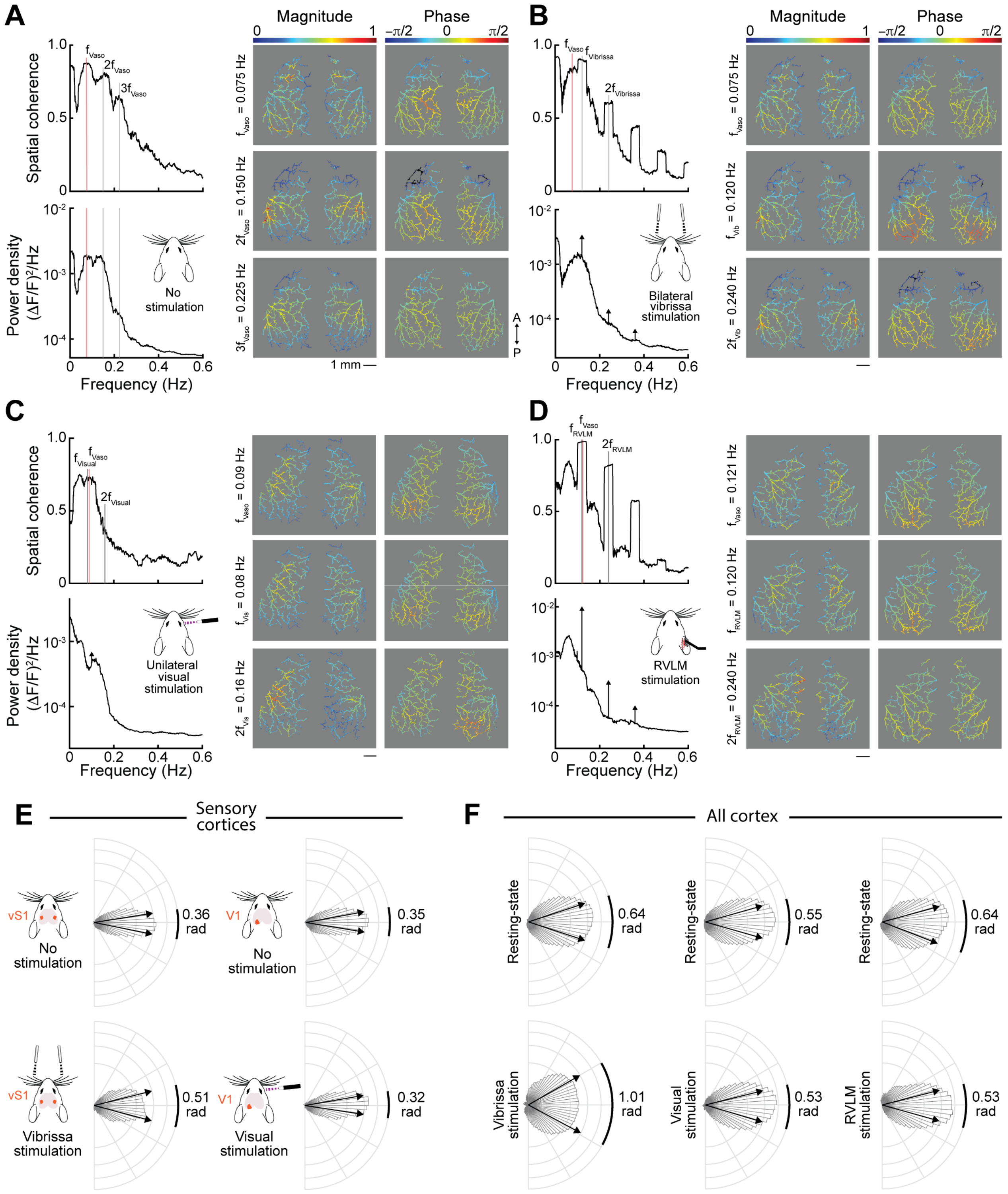

Figure S8. Duckworth*, Broggini*, Chhabria, Neill, Sancho, Sellier-Prono, Iyengar, Nelson, Vergassola & Kleinfeld

**Figure S7: Pial arteriole spatial coherence and propagation patterns during resting-state and stimulated conditions**

**(A)** Coherent resting-state activity at the peak vasomotor frequency and higher harmonics for one example experiment. Magnitude and phase maps are calculated from the leading mode of a space-frequency SVD of the data (**STAR Methods**). Temporal delay increases with increasing phase.

**(B)** Coherent activity patterns during an example bilateral vibrissa stimulation experiment. Frequencies of interest were chosen as the resting-state vasomotor frequency, stimulation frequency, and stimulation frequency 2$^{nd}$ harmonic.

**(C)** Coherent activity patterns during an example unilateral visual stimulation experiment. Plotted as in (**B**).

**(D)** Coherent activity during an example RVLM activation experiment. Plotted as in (**B**).

**(E)** Distribution of phase values within vibrissa and visual primary sensory cortices during rest and stimulation. Distributions include data summarized across experiments within each dataset (i.e. visual or vibrissa stimulation datasets). Each experiment's phase distribution was centered around its circular mean before being added to the plot. Only experiments with $0.095 < f_{stim} < 0.135$, or $0.095 < f_{vaso} < 0.135$ for resting-state experiments, were used, as vasomotor phase gradients vary with frequency[1]. Arrows and bold lines denote the central 50$^{th}$ percentile.

**(F)** Phase distributions across the imaged cortical mantle during sensory and RVLM stimulus experiments, plotted as in (**E**). Columns are common datasets.

1. Broggini, T., Duckworth, J., Ji, X., Liu, R., Xia, X., Mächler, P., Shaked, I., Munting, L.P., Iyengar, S., Kotlikoff, M., et al. (2024). Long-wavelength traveling waves of vasomotion modulate the perfusion of cortex. Neuron *112*, 2349-2367.
